# Discovering and exploiting active sensing motifs for estimation with empirical observability

**DOI:** 10.1101/2024.11.04.621976

**Authors:** Benjamin Cellini, Burak Boyacıoğlu, Stanley David Stupski, Floris van Breugel

## Abstract

Organisms and machines must use measured sensory cues to estimate unknown information about themselves or their environment. Cleverly applied sensor motion can be exploited to enrich the quality of sensory data and improve estimation. However, a major barrier to modeling such active sensing problems is the lack of empirical, yet rigorous, tools for quantifying the relationship between movement and estimation performance. Here, we introduce “BOUNDS: Bounding Observability for Uncertain Nonlinear Dynamic Systems”. BOUNDS can discover patterns of sensor motion that increase information and reduce uncertainty in either real or simulated data. Crucially, it is suitable for high dimensional and partially observable nonlinear systems with sensor noise. We demonstrate BOUNDS through a case study on how flying insects estimate wind properties, showing that specific active sensing motifs improve estimation. Additionally, we present a framework to refine sporadic estimates from active sensing. When combined with an artificial neural network, we show that the information gained via active sensing in real *Drosophila* flight trajectories is suitable for precise wind direction estimation. Collectively, our work will help decode active sensing in organisms and inform the design of estimation algorithms for machines.

## INTRODUCTION

A fundamental challenge for any agent—organisms and robotic systems alike—is to combine and transform incoming sensory cues into estimates of egocentric or allocentric task-relevant variables. Often, available sensory cues do not directly measure these variables, but encode them indirectly. For example, orienting towards an object in the field of view first requires decoding visual information from the retina (or pixels from a camera) to infer the object’s location ^1,2^. For a moving agent, sensory cues will be dynamic, and inferring task relevant variables often requires integrating measurements over some time window. For instance, estimating object velocity from position measurements requires comparing measurements over time to estimate a derivative. In many cases, however, variables of interest are not linearly related to available sensor measurements, or their derivatives.

For an agent with nonlinear sensor or movement dynamics, different types of motion can help to decouple correlated variables. This means that intelligent sensor motion can potentially enhance an agent’s ability to perform an estimation task, a process we refer to as *active sensing* ^3^. For example, many organisms extract image angular velocity, referred to as optic flow, from their visual system to encode their movement relative to nearby objects. Optic flow captures the ratio of velocity and distance, making it difficult to estimate either one independently. However, by changing velocity with a known acceleration these two variables can be decoupled from optic flow to estimate both distance ^4^ and velocity ^5^. In this example, acceleration allows a monocular agent to capture similar information that would be available to a stationary agent with two separate imaging sensors with binocular overlap and known geometry. In this sense, active sensing can help make up for a lack of sensors, or provide redundant estimation pathways.

Many organisms perform movements, either of their whole body or of individual sensors, that might serve some active sensing role. For example, mantids perform peering head movements before striking prey ^6^, a behavior thought to provide them with distance information. Mice also make similar head movements before jumping over a gap ^7^. Rats ^8^ and nectar feeding moths ^9^ actively move their whiskers or proboscis across objects to infer 3-dimensional properties. Whereas in these examples active sensing is thought to help animals estimate otherwise inaccessible variables, sensor movement can also simply improve the quality of information. For example, head stabilization reflexes in flies ^10,11^ and birds ^12^, pinna movements of bats ^13^, and active antenna positioning in flies ^14^ are all thought to maximize the signal to noise ratio. How can we understand what extra information is being acquired via body or sensor movements while considering the time-history of all putative sensory measurements? In simple terms, why is movement beneficial in a sensory context? A computational approach for rigorously discovering active sensing motifs would serve as a powerful hypothesis generating tool to help explain many comparable sensorimotor behaviors across taxa, while simultaneously helping engineers to optimize estimation for autonomous machines.

Recent approaches for quantifying the information made available through active sensing in organisms leverage *observability* ^15,16^. Observability is a control-theoretic concept that describes which states can be estimated from a set of sensor measurements and associated dynamics model ^17^. States are variables of interest that capture the internal and external factors that influence a system’s behavior. In the context of biological systems, states could be internal variables such as an organism’s position, velocity, or heading—or could be external variables like a prey/predator position or an environmental quantity (e.g. wind, terrain, etc.). It is useful to be able to model which states can, or can not, be estimated from an agents available sensory cues because it can reveal the potential goal of sensorimotor behaviors. For instance, when seeking refuge under low luminance, weakly electric fish increase oscillations in their swimming patterns ^18,19^. This behavior is predicted by a loss of observability of the fish’s position in a refuge due to high-pass sensor dynamics ^16,20^. Whereas existing observability approaches can provide insight into these types of problems, established methods are poorly suited for high dimensional and/or partially observable nonlinear systems, do not offer a physical unit for observability measures, and do not consider the impact of sensor noise. Furthermore, there are no established approaches for applying observability tools to experimental data.

Here we introduce a novel computational pipeline— “BOUNDS: Bounding Observability for Uncertain Dynamic Systems”—that leverages tools from control and information theory to address these shortcomings. We extend established methods for empirically calculating an observability matrix (𝒪) for nonlinear systems ^15,21^ to work with real data through a model predictive control framework. To take sensor noise properties into account, we calculate the Fisher information matrix (ℱ) associated with 𝒪^22^. We introduce a regularization approach that allows us to invert ℱ under rank-deficient conditions so that we can apply the Cramér–Rao bound (CRB) ^23^ on partially observable systems to quantify observability with meaningful units. Iterating our pipeline along a dynamic trajectory intuitively reveals active sensing motifs. Our approach can answer questions pertaining to sensing in biological and robotic systems such as:

1. What can be estimated?
2. How accurate can an estimate be?
3. What sensors are required?
4. What kind of active sensor motion is required or helpful?

Finally, we present a framework for leveraging observability information—or the movement motifs correlated with observability—to filter and refine estimates. This approach— “MIF: Motif Informed Filtering”—may allow organisms and machines to stitch together sporadic estimates corresponding to bouts of active sensing.

## RESULTS

To describe our computational pipeline, as well as new scientific discoveries, we have organized this section as follows. First, we provide a step-by-step description of our computational pipeline using Figure 1 as a visual organization of these steps. Second, we introduce a specific case study— anemosensing (sensing properties of ambient wind) in flying insects—to showcase the different ways our method can be used to discover active sensing motifs from simulations and data. Finally, we introduce a framework for incorporating knowledge of observability—or motifs correlated with observability—into an estimator to provide continuous estimates across sporadic active sensing bouts, and again illustrate this approach using our anemosensing case study.

**Figure 1.**
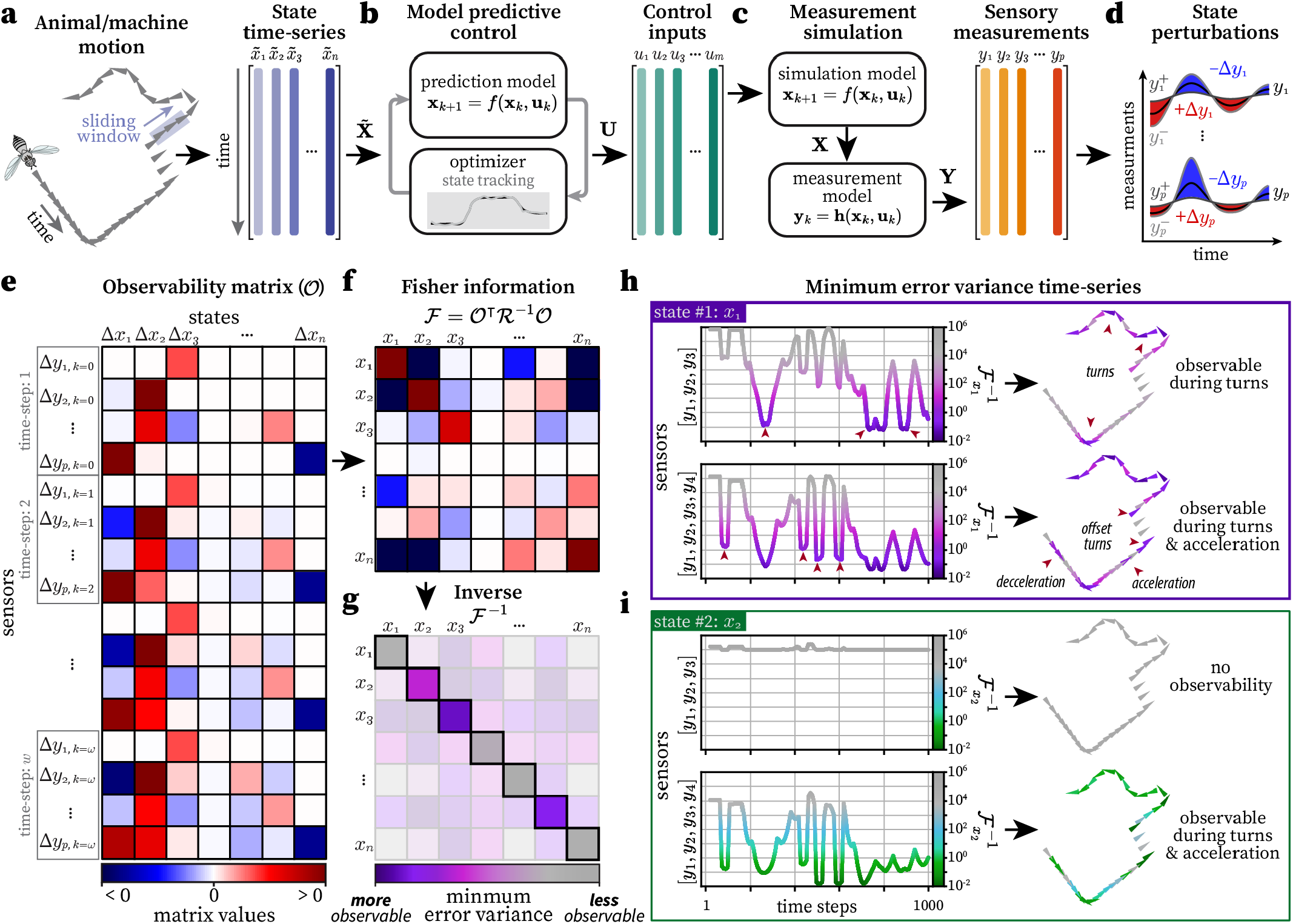
Overview of BOUNDS, a method for discovering active sensing motifs with empirical observability. **a**. A simulated or measured state trajectory (in this case of a flying insect) that is described by a collection of time-series of internal and external state variables 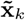. **b**. Model predictive control is used to find the control inputs **u**_*k*_ required to reconstruct the state trajectory in *a* through a simulation model (Eq. 11). **b**. The MPC control inputs **u**_*k*_ are used to simulate the dynamics associated with the state trajectory 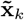. The simulated state trajectory **x**_*k*_ is used to with a measurement model **h**(**x**_*k*_, **u**_*k*_) to simulate the sensory measurements. **d**. Each initial state *x*_*i*,0_ is perturbed in the positive *x*_*i*,0_^+^ and negative *x*_*i*,0_^−^ directions. Then the corresponding change in sensor measurements over time, **y**_*k*_^+^ and **y**_*k*_^−^, are computed. **e**. An illustration of an observability matrix 𝒪, where the color grid indicates the values in the matrix (corresponding to *c*). The columns indicate the difference of the initial state perturbation Δ*x*_*i*,0_ and the rows indicate the corresponding difference in measurements Δ**y**_*k*_ over time for a discrete number of time steps *ω*. **f**. An illustration of the Fisher information matrix, calculated as ℱ= 𝒪^⊺^R^−1^ 𝒪, where R is the noise covariance matrix. Colors indicate matrix values as in *e*. **g**. The inverse of the Fisher information matrix ℱ^−1^, which encodes the minimum error covariance of the estimate of the state’s initial condition **x**_0_. The diagonal elements represent the minimum error variance of each individual state—the inverse of the observability level—where smaller values indicate a higher observability level. **h**. The minimum error variance time-series (arbitrary units) of the first state *x*_1_ pulled from ℱ^−1^, where the color corresponds to the observability along the trajectory in *a*. Turns are observable for the first three sensors [*y*_1_, *y*_2_, *y*_3_] (top) but adding a fourth sensor *y*_4_ also renders acceleration/deceleration and turns offset to heading observable. The color-map here is the same as in *f*, because the error variance is pulled from ℱ^−1^ in sliding windows. **i**. Same as *g* but for the second state *x*_2_. Note that *x*_2_ is completely unobservable for sensors [*y*_1_, *y*_2_, *y*_3_] but adding *y*_4_ yields similar observability to *x*_1_ in *h*. The color-map here is distinct from *f* to highlight that a different state is being considered, but the error variance is still pulled from ℱ^−1^ in *f*.

### A method for discovering active sensing motifs with empirical observability

#### 1. Determine states and collect state trajectory time-series

Consider a discrete time state trajectory 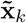 that describes the movement of an agent or sensor, as well as environmental variables (Figure 1a). The primary objective is to compute a new time-series that describes the observability of a given state variable(s) along this trajectory, which can be used to evaluate the efficacy of putative active sensing motifs. All relevant state variables that describe the behavior and environment of the agent must be included. The state trajectory can be simulated for testing hypotheses, or measured from real data to evaluate the observability properties of an agent’s true locomotion patterns.

#### 2. Define a model

Once a collection of state variables is determined, a model of the agent’s dynamics must be defined. In general, a good model accounts for 1) the inputs that control locomotion (e.g, forces and torques) 2) basic physical properties such as mass, inertia, damping, etc., and 3) sensory dynamics that model what an agent is measuring (Figure 1b). We describe our approach for a system with nonlinear deterministic dynamics, and stochastic measurements as

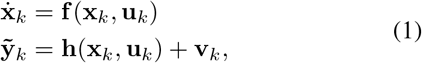

where **x**_*k*_ is the state vector representing *n* states over time, **u**_*k*_ is the input vector, and 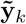 is the measurement vector corresponding to *p* measurements at time step *k*. The function **f** (**x**_*k*_, **u**_*k*_) describes the system dynamics. The function **h**(**x**_*k*_, **u**_*k*_) describes the measurement dynamics, and **v**_*k*_ is some additive Gaussian noise with covariance matrix R_*k*_. The effect of the noise on observability is taken care of in step 6. Steps 3-5 only use the deterministic measurement equation **y**_*k*_ = **h**(**x**_*k*_, **u**_*k*_). A benefit of our empirical approach is that the model does not have to be closed-form, and can instead be a simulation or computational model ^24,25^—for instance, a physics engine model of *Drosophila* ^26^. The only requirement is the ability to simulate the system forwards in time. The accuracy of the resulting observability analysis will depend on the faithfulness of the model to true physiology and biomechanics.

#### 3. Reconstruct state trajectory through model predictive control

Next, our method applies model predictive control (MPC) to find the inputs **u**_*k*_ required to reconstruct the state trajectory with the chosen model ^27^ (Figure 1b). This step is critical for nonlinear or time-varying models where observability can change as a function of the state variables and inputs. The MPC determined inputs are used to simulate the nominal trajectory **x**_*k*_ (which should be very similar to the observed trajectory 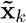) and associated measurements **y**_*k*_ (Figure 1c). Although other control methods could suffice, properly tuned MPC will typically yield the closest match between the simulated and desired trajectory.

#### 4. Quantify measurement sensitivity to the initial state

With a model and inputs in hand, the next objective is to quantify the sensitivity of the sensory measurements **y**_*k*_ along the state trajectory relative to the nominal initial state **x**_0_ (Figure 1d). Note that observability quantifies how feasible it is to estimate the *initial state* of a system given future measurements from some time window of length *ω*. The dual concept of constructability can be used to analyze the final state ^28^, but requires either discretizing the dynamics or inverting the continuous time dynamics. Here we focus our analysis on observability due to its relative computational efficiency, but note that for small time windows the two will generally be strongly correlated. The sensitivity of the sensor measurements to small changes in the initial state can be compactly represented as the Jacobian of the measurements over time with respect to the initial state, Δ**Y**_*ω*_*/*Δ**x**_0_. Here we use **Y**_*ω*_ to denote the collection of *p* measurements for each time step from *k* = 0 : *w*. Following established methods ^24,25^, we empirically approximate the Jacobian by applying small perturbations to each initial state *x*_*i*,0_, one at a time, in the positive 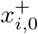 and negative 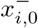 directions, simulate the system using the inputs **u**(*t*) calculated in step 3 (they are not recalculated for the perturbed trajectory), and then compute the difference in the corresponding change in sensory measurements across time 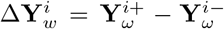 (Figure 1c). The end result for each state perturbation is a column vector of length *p × ω*, corresponding to *p* measurements over *ω* time steps.

#### 5. Construct observability matrix

Repeating step 4 for each state variable and stacking the resulting column vectors side by side yields the full empirically determined Jacobian, Δ**Y**_*ω*_*/*Δ**x**_0_, which is equivalent to the empirical observability matrix 𝒪 (Figure 1e). The columns of 𝒪 indicate perturbations to each initial state, whereas the rows indicate the corresponding change in sensory measurements for each measurement and time step. A state variable is observable if the rows of 𝒪 can be combined to reconstruct the basis vector corresponding to that state variable ^15^. However, quantifying the *level* of observability for each state variable is more challenging. Prior approaches have used metrics to measure the level of observability such as a rank-truncated condition number for a subset of the rows of 𝒪^15^ or the eigen-values of the observability Gramian ^25^. These metrics are either numerically fragile ^15^ or do not work on partially observable systems ^25^, and neither account for sensor noise.

#### 6. Compute the Fisher information matrix and it’s inverse

To compute observability levels for individual state variables we construct the Fisher information matrix as

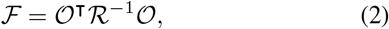

where ℛ is a block diagonal matrix of the measurement noise covariance matrices at each time step. For the deterministic dynamics we consider, the process noise *Q* is 0 (see Methods) (Figure 1f) ^29,30^. The Fisher information matrix captures the amount of information that a known random variable, in our case the measurements **Y**_*w*_, carries about unknown parameters, in our case the initial state **x**_0_. The Cramér–Rao bound states that the inverse of the Fisher information matrix ℱ^−1^ sets the lower bound on the error of the state estimate ^23^. Thus, the diagonal elements of ℱ^−1^ tell us the minimum possible variance in the error for each state variable (for any estimator), inversely correlating with observability (Figure 1g).

In cases where there is at least one unobservable state, ℱ will not be invertible. Therefore, to compute ℱ^−1^ we add a small regularization term to ensure invertibility ^31^ with minimal impact on the minimum error covariance:

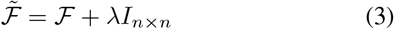

where

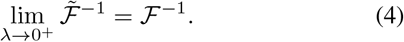

For unobservable states, this “Chernoff inverse” yields large values in the corresponding diagonal elements, whereas the more traditional Moore-Penrose pseudo-inverse yields misleading values. For real values of *λ* this procedure does place an upper bound on the minimum error covariance, and thus *λI*_*n×n*_ must be chosen carefully, so as to not inject too much “artificial” observability into the system, while still allowing an inverse to be computed (see Methods).

#### 7. Reveal active sensing motifs through sliding windows

Finally, we repeat steps 1-6 in sliding time windows, that is, we use the same window size *w* but with initial conditions that start progressively later in time along the nominal trajectory. For each window we construct 𝒪 and ℱ and calculate the minimum error variance for each state variable from 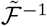. This yields a time-series of minimum error variance along the state trajectory (Figure 1h–Figure 1i). Because observability analyzes the initial state given future measurements, our metric would show high observability levels immediately *before* an active sensing maneuver. To make the relationship between temporal patterns in observability and features of the trajectory more intuitive, we shift the time series back in time by half of the sliding window length, *ω/*2. Note that it is possible to rigorously evaluate the total information that corresponds to the middle of a time window given both the past, present, and future measurements, but this comes at a much higher computational expense ^28^.

### 8. Approach summary

Our approach allows us to evaluate the temporal patterns of observability as a function of the state variables and available sensory measurements. For instance, in a toy example for the state trajectory in Figure 1a, we can see that changes in heading angle lead to an order of magnitude decrease in the minimum error variance—analogous to an increase in observability—for the first state 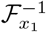, compared to a straight trajectory (Figure 1h). This result is based on three sensory measurements [*y*_1_, *y*_2_, *y*_3_], however adding a fourth sensor *y*_4_ also renders acceleration/deceleration and turns offset to the heading angle more observable (Figure 1i). This illustrates that active sensing motifs can be tuned to the sensory data that is available. A critical point is that each state can have different observability properties, and thus distinct active sensing motifs. For example, in contrast to *x*_1_, the second state *x*_2_ is never observable with just [*y*_1_, *y*_2_, *y*_3_], but adding *y*_4_ leads to similar active sensing patterns seen for *x*_1_ (Figure 1h). This approach allows us to investigate fundamental relationships between states, sensors, and active sensor motion.

### Case study: discovering active sensing motifs for anemosensing

To illustrate the opportunities for insight that our approach provides, we applied it to an open question in active sensing: how do flying insects estimate properties of ambient wind (anemosensing)? This is a critical component of olfactory navigation, and behavioral evidence suggests that insects do estimate wind direction (and to some extent magnitude) in flight ^32^. However, insects cannot measure ambient wind 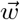 directly. Instead, insects use deflections of their antennae ^33,14^ to measure *apparent* airflow 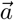, the vector sum of the ambient wind 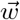 and motion-induced airflow 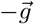, where 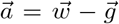 (Figure 2a). If insects could measure their ground velocity vector 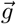 in addition to 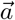, both in consistent and calibrated units, it would be trivial to perform a vector summation to solve for the ambient wind 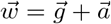. Since insects cannot measure 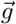 directly, however, they must estimate 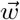 through some other approach.

**Figure 2.**
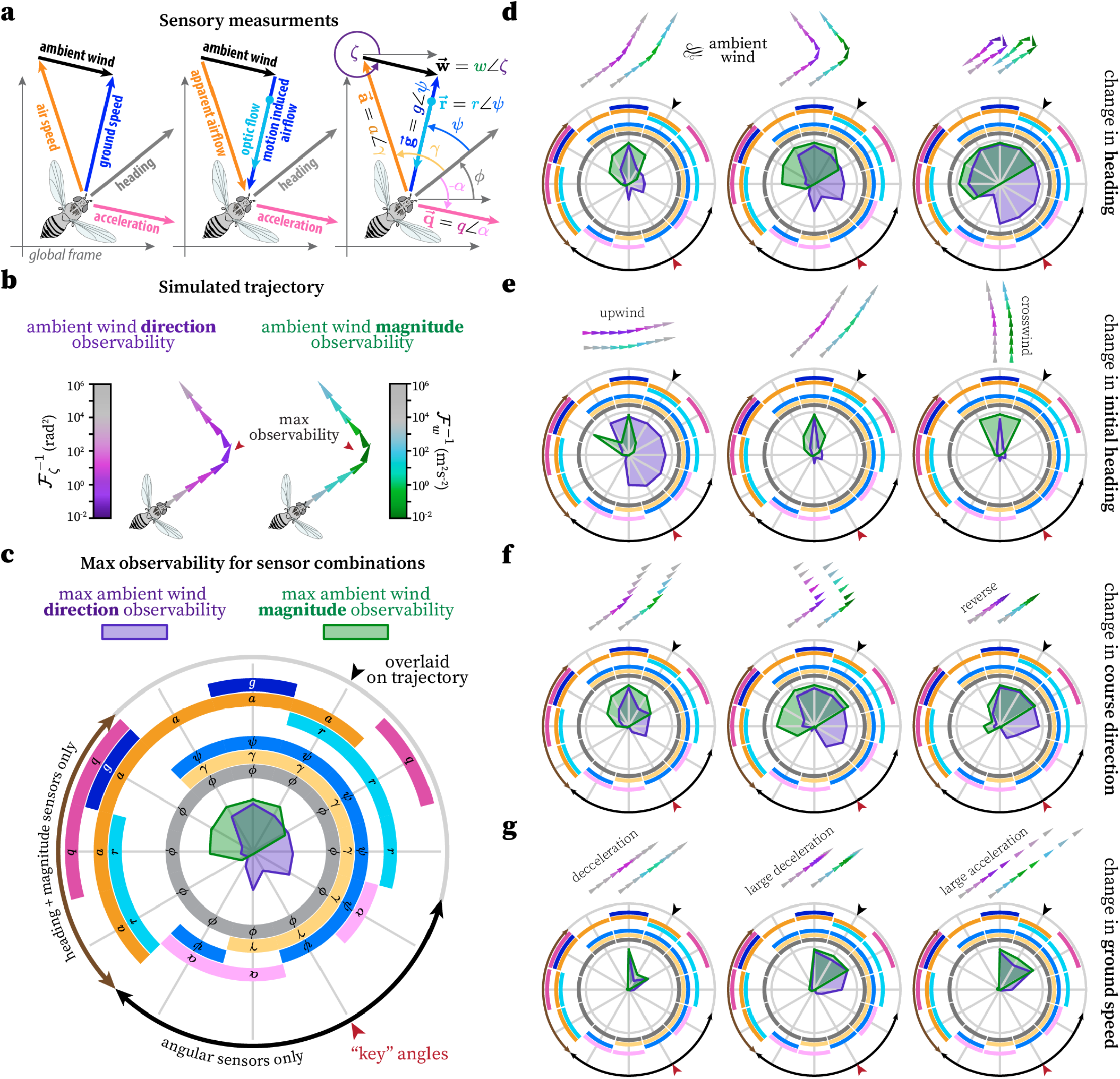
Discovering active sensing motifs for anemosensing. **a**. Illustration of the trigonometric properties of an agent in the presence of ambient wind (left) and the corresponding putative sensory measurements (middle). Each vector quantity is decomposed into a magnitude and angle (right). **b**. An example simulated flight trajectory with the level of observability (minimum error variance) of ambient wind direction (purples) and magnitude (greens) indicated by the color. Observability was calculated for a 40 ms time window, corresponding to 4 discrete measurements (*ω* = 4). We use this same time window for all subsequent analyses. **c**. Radar plot illustrating the maximum observability (normalized and inverted minimum error variance) of ambient wind direction (purple) and magnitude (green) of the trajectory from *b* for many combinations of sensors. The circular grid indicates which combination of sensors were considered and the radii of the patch pointing towards at each grid line indicates the observability level of the corresponding sensor set. Note that each sensor has a corresponding color: heading (gray), apparent airflow (orange), ground speed (blue), acceleration (pink) corresponding to *a*, where magnitudes are dark colors and angles are light colors. Optic flow magnitude (cyan) is associated with the same angle as ground velocity. The black arrowhead on the outside of the grid indicates the sensor combination whose observability is overlaid on the trajectory. **d**. Radar plots for simulated trajectories with turns of varying magnitude. Ambient wind direction (coming from the right) and magnitude were set to be constant (see Methods for full simulation details). Reference *c* for sensor legend. **e**. Same as *d* but varying the initial heading of the trajectory, and a constant small turn magnitude. Note that passing through or near upwind dramatically increases the observability of wind direction. **f**. Same as *d* but for turning with out changing heading (pure change in course direction). **g**. Same as *d* but for a change in ground speed (acceleration/deceleration).

What sensory information do insects have available? Much of the navigational hub of the insect brain, the central complex, primarily encodes the angular quantities associated with apparent airflow *γ* and ground velocity *ψ* ^34,35,36^, all relative to an estimate of heading *ϕ* in world coordinates ^37^. With just these three angles, wind direction is observable, but only during a turn ^38^. Magnitude information is also present, however, it is unlikely to be in consistent and calibrated measurement units. For example, instead of ground speed *g* insects have access to optic flow cues 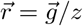, which depend on elevation *z*, and their encoding can be a function of texture ^39^ and behavioral state ^40^. How the inclusion of the magnitudes *r* and *a* with various angular components affect observability is not well understood. This scenario is akin to computing all the angles and sides of a triangle given only one angle and one side. The only way to resolve this under defined problem is through temporal integration of sensory information during an active sensing maneuver.

### Evaluating putative active sensing motifs

To investigate which putative active sensing strategies insects could employ to estimate ambient wind speed and direction, we simulated a variety of flight trajectories with distinct motifs, such as turns and accelerations (Figure 2). We then applied our observability analysis in sliding windows along each trajectory and extracted the maximum observability (minimum error variance) for both the ambient wind direction and magnitude (Figure 2b).

We considered a variety of possible sensor groups consisting of combinations drawn from four angular sensors: heading *ϕ*, ground velocity angle *ψ*, apparent airflow angle *γ*, and acceleration angle *α* (Figure 2a), and four magnitude sensors: ground speed *g*, apparent airflow magnitude *a*, acceleration magnitude *q*, and optic flow magnitude *r* = *g/z*, where *z* is the distance above the ground. We do not know which sensors out of these eight flies might use for wind estimation, so we examined some of the most likely combinations (out of 255 possibilities). We paid particular attention to three “key angles” identified in the literature ^38,41^ **y** = [*ϕ, ψ, γ*] (Figure 2, red arrows) with small variations thereof, such as a combination that includes both airflow and optic flow magnitude, **y** = [*ϕ, ψ, γ, a, r*] (Figure 2, black arrows). As a reference, we also considered two biologically implausible combinations that include ground speed *g*. As expected from a vector subtraction operation, the 12-o-clock combination of [*ϕ, ψ, γ, a, g*] is always observable. We first considered a pure turning motif with constant ground speed and no side-slip. Consistent with prior work ^38^, we found that turning was necessary for estimating wind direction for angular combinations such as **y** = [*ϕ, ψ, γ*] (Figure 2d, *purple*). Our analysis extends this result to other combinations, such as those with optic flow or airflow magnitude. In general, larger turns (*>* 90^°^) were required for high observability. Interestingly, wind direction was highly observable for the largest turns for many sensor combinations of just angular sensors, implying that the wind direction computation may not need to rely on magnitude information (Figure 2d). However, at least one magnitude sensor that included a distance unit (e.g., meters)—ground speed, apparent airflow, or acceleration, but not optic flow—was necessary for the wind magnitude to be observable (Figure 2d). Wind magnitude, but not wind direction, was also generally observable if only heading and magnitude sensory cues were available (Figure 2d). This suggests that there could be distinct neural implementations for estimating wind direction and magnitude that use different sensory cues, even if the active sensing motif (turning) is the same.

Other factors such as the absolute heading of a trajectory, as opposed to a *change* in heading, also affect the observability of the wind. Trajectories that start near, or pass through, the upwind (0^°^) or downwind (180^°^) direction exhibit much higher observability of wind direction, even for small turns (Figure 2e). These trajectories pass through either the stable or unstable fixed points for the visual anemotaxis strategy, which uniquely identifies wind direction ^41^. Therefore, a generally effective active anemosensing strategy is to perform a sufficiently large turn that is likely to pass through upwind/downwind. Furthermore, once aligned upwind, maintaining an upwind course should be possible without performing additional large turns. Passing through upwind/downwind as a strategy is not helpful for estimating wind magnitude, however orienting crosswind is beneficial for a select sensor set with the three key angles and apparent airflow magnitude **y** = [*ϕ, ψ, γ, a*] (Figure 2e).

Although flying insects often fly with their heading aligned with their course direction, as in (Figure 2d), this is not always the case as some flies have been observed traveling nearly perpendicularly to their own heading ^42^. Therefore, we investigated an analogous active sensing motif consisting of an offset turn (Figure 2f). Overall, this motif was similar to turning with a change in heading, but exhibited higher wind direction observability for the key **y** = [*ϕ, ψ, γ*] angular sensor set (Figure 2f). Wind direction was unobservable without *ψ*, which represents the direction of travel with respect to heading. It is intuitive that *ψ* is critical for this motif because there there is no change in heading *ϕ*, and all of the rich temporal information from the turn is contained within the *ψ* sensory cue. The special case of a 180^°^ turn displayed lower observability. This maneuver corresponds to a reversal of course direction.

Flying insects often modulate their ground speed, especially in wind ^43^ and while tracking odors ^32^. We investigated how changes in ground speed—i.e., acceleration or deceleration— impacted observability. We found that both large acceleration and deceleration lead to high observability of wind direction and magnitude. In contrast with turning motifs, changes in ground speed require at least one magnitude measurement for wind direction be observable (Figure 2g). For estimating wind direction, optic flow magnitude was sufficient as a magnitude measurement, whereas wind speed required a measurement of acceleration or airflow magnitude. At least some of the observability associated with deceleration may come from the overall change in ground speed relative to the wind speed. A low ground speed relative to the wind speed results in more observable wind direction (Figure S1). The opposite is true for a high ground speed relative to the wind speed, for which the wind direction is less observable, but the wind magnitude is much more observable (Figure S1). This could explain why flies can readily detect the presence or absence of wind ^32^.

### Effects of dynamic wind during anemosensing

The previous section considered how an agent’s active changes in course or speed affect observability. Movement can also be influenced by external forces. To illustrate how external perturbations can influence observability we explored how dynamic wind affects a flying insects ability to estimate wind direction and magnitude. We simulated flight trajectories during wind gusts (increase in wind magnitude) and changes in wind direction. We considered two cases, 1) where the agent does not react to the change in wind and is blown off course (Figure 3a) and 2) where the agent maintains course by applying controls to reject wind induced motion (Figure 3b).

**Figure 3.**
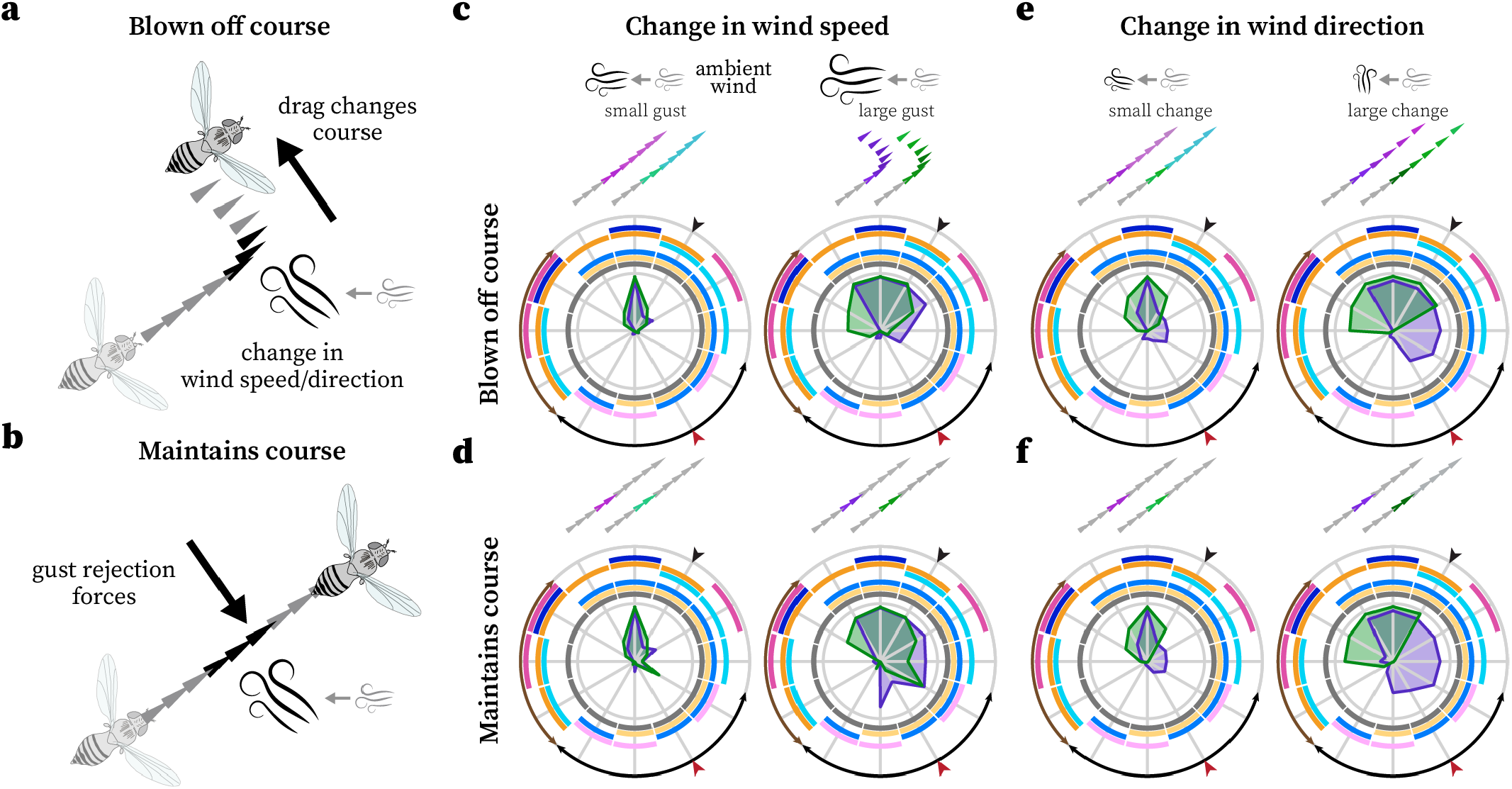
Dynamic wind can make wind properties observable. **a**. Simulation of a change in wind magnitude and/or direction blowing an agent off course via drag forces, where the agent does not attempt to reject the change to its course. **b**. Same as *a* but the agent applies flight forces to reject the perturbation to its flight path and maintain course. **c**. The observability of wind direction (purple) and magnitude (green) for different combinations of sensors for wind gusts (change in wind magnitude) of varying magnitude, without rejection forces. Refer to Figure 2 for legend. **d**. Same as *c* but with rejection forces to maintain course. **e**. Same as *c* but for a change in wind direction. **f**. Same as *d* but for a change in wind direction.

We found that wind gusts large enough to change an agent’s course lead to high observability of wind direction and magnitude but, similar to the acceleration motif, require at least one magnitude measurement (Figure 3c). This suggests that gusts alone may provide enough information for flying insects to orient into the wind, without requiring an active sensing motif like in constant wind (Figure 2). In the case where an agent maintains course during a gust we found that angular sensor groups including the acceleration angle *α* were significantly more observable than when the agent is blown off course by a gust (Figure 3d). This is likely because *α* provides useful information about the flight force direction required to reject the gust.

Observability during changes in wind direction followed a similar trend to wind gusts: large changes in wind direction increased observability. However, some of the angular sensor sets displayed high wind direction observability without requiring a magnitude measurement, both when the agent maintained, and was blown off, course. (Figure 3e–Figure 3f). The acceleration angle *α* was still beneficial, but not required, when the agent maintained course (Figure 3f). Interestingly, some sensor sets only containing magnitude measurements and the heading showed high wind speed observability, even though only the wind direction changed (Figure 3e–Figure 3f). Altogether, changes in the wind direction and magnitude may be informative for flying insects.

### Methodical discovery of active sensing motifs

Thus far, we only investigated discrete motifs (pure turning, acceleration, etc.) that were inspired by prior work in the literature. Now we illustrate how active sensing motifs can be discovered using a more agnostic, methodical approach which can also consider combinations of discrete motifs. We simulated trajectories within a three dimensional state and action space consisting of combinations of initial heading, angular velocity, and acceleration/deceleration. We computed the observability of wind direction for each unique trajectory for many sensor groups and examined the effects of blending motifs.

We first verified that that the pure active sensing motifs we discovered (Figure 2) were captured by this approach. This was the case, as pure turning resulted in increased observability, crossing through the upwind direction (0^°^) was beneficial even for small turns, and acceleration alone only had an effect for sensor groups that include a magnitude (Figure 4). The most observable trajectories were generally those that started with an initial heading offset from upwind (±90^°^ or ±45^°^), made a large turn crossing through upwind, and also decelerated (Figure 4). This result demonstrates that there can be a benefit to blending active sensing motifs, which was not clear from our prior analysis. This interaction was most pronounced for turns away from upwind, where adding deceleration led to a much larger increase in wind direction observability (Figure 4). Surprisingly, positive acceleration was sometimes detrimental to wind direction observability (Figure 4), likely because of the resulting increase in the ground speed to wind speed ratio (Figure S1). Accelerating was still better than flying straight at constant velocity, but for a trajectory that is already turning, adding deceleration as opposed to accelerating is a better option. This indicates that blending motifs to optimize observability requires a careful analysis.

**Figure 4.**
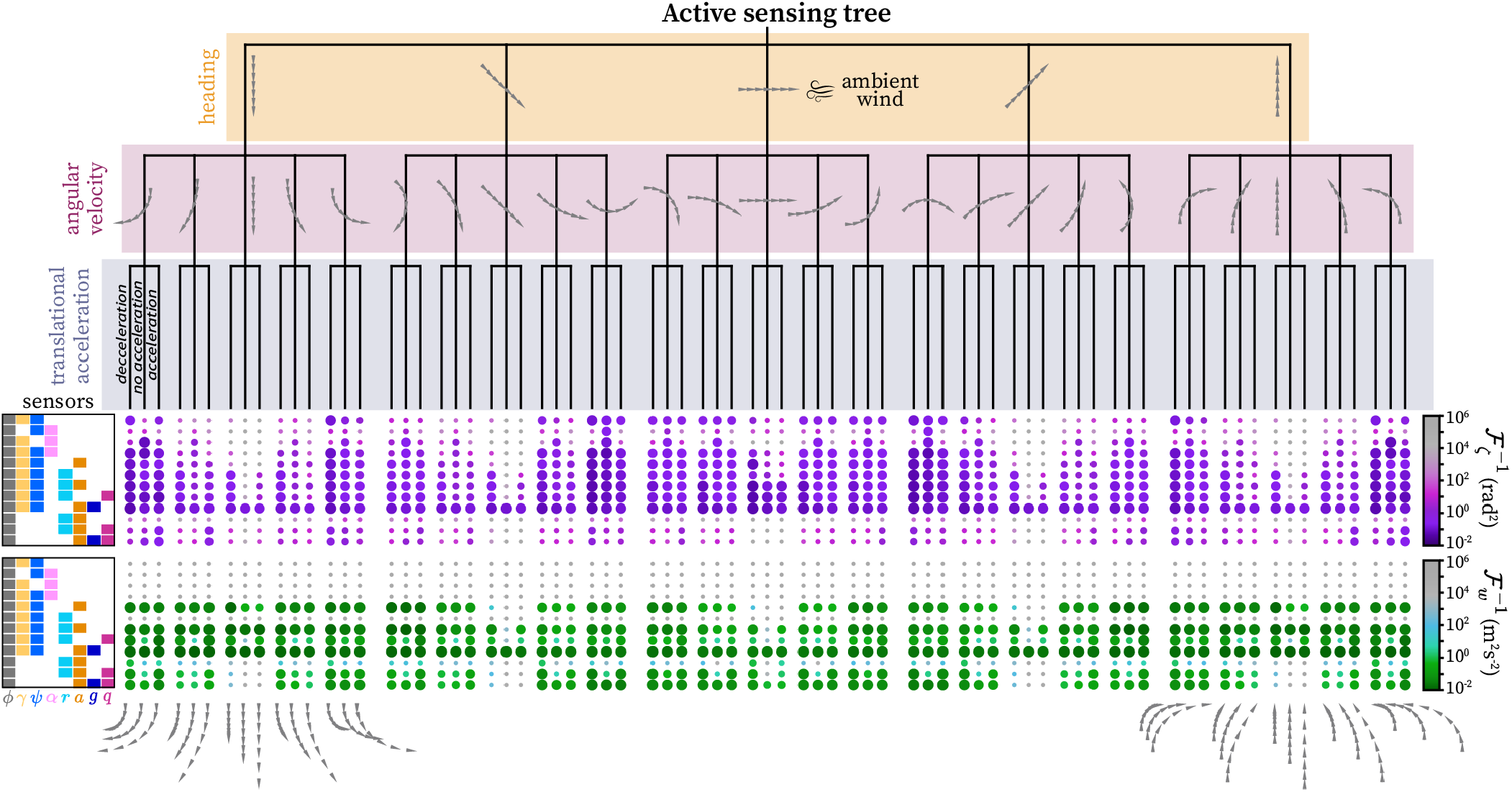
Methodical discovery of active sensing motifs. A tree representing various combinations of active sensing motifs and the corresponding observability of wind direction (purple nodes) and wind magnitudes (greens nodes). The trajectories were created by starting with a given heading (top branch), adding angular velocity (middle branch), and then adding acceleration/deceleration (bottom branch). Each trajectory for the heading and angular velocity branches are overlaid on the tree, but the acceleration branch trajectories are only shown for the left and right foremost branches for visual clarity. The observability of each unique trajectory for various sensor groupings are represented by the size and color of the nodes at the end of the leaves, where larger nodes are more observable. The legend on the left indicates which sensor groupings were used for the observability calculation (based on Figure 2 variables and colors).

### Observability properties of anemosensing behavior in measured fly flight

An important improvement of our approach over prior observability tools is that it can be applied to real data by using model predictive control to drive a system model along data from an observed trajectory. Here we illustrate this feature using previously published trajectories of freely flying *Drosophila* engaged in anemosensing ^32^ to explicitly answer the question of whether flying insects move in a manner that enhances information gathering potential. In brief, flies were placed in a wind tunnel with constant 0.4 m s^−1^ wind and allowed to fly freely. To achieve high spatiotemporal control over their olfactory experience and corresponding wind-orienting behavior, an optogenetic approach was used to deliver a 675 ms fictive olfactory experience by flashing red light to activate Orco^+^ neurons expressing CsChrimson, a red-light-sensitive ion channel (for specific method details see Stupski and van Breugel ^32^) (Figure 5a). The fictive odor experience evoked a strong upwind orienting response but, as highlighted in Stupski and van Breugel ^32^, flies accomplished this with a rapid sequence of two distinct turns (Figure 5a–Figure 5b). The first turn was not directed upwind ^32^, instead ending in a random direction (Rayleigh z-test, *Z* = 0.0, *p* = 0.66) (Figure 5b), supporting the notion that this turn may serve an information gathering role.

**Figure 5.**
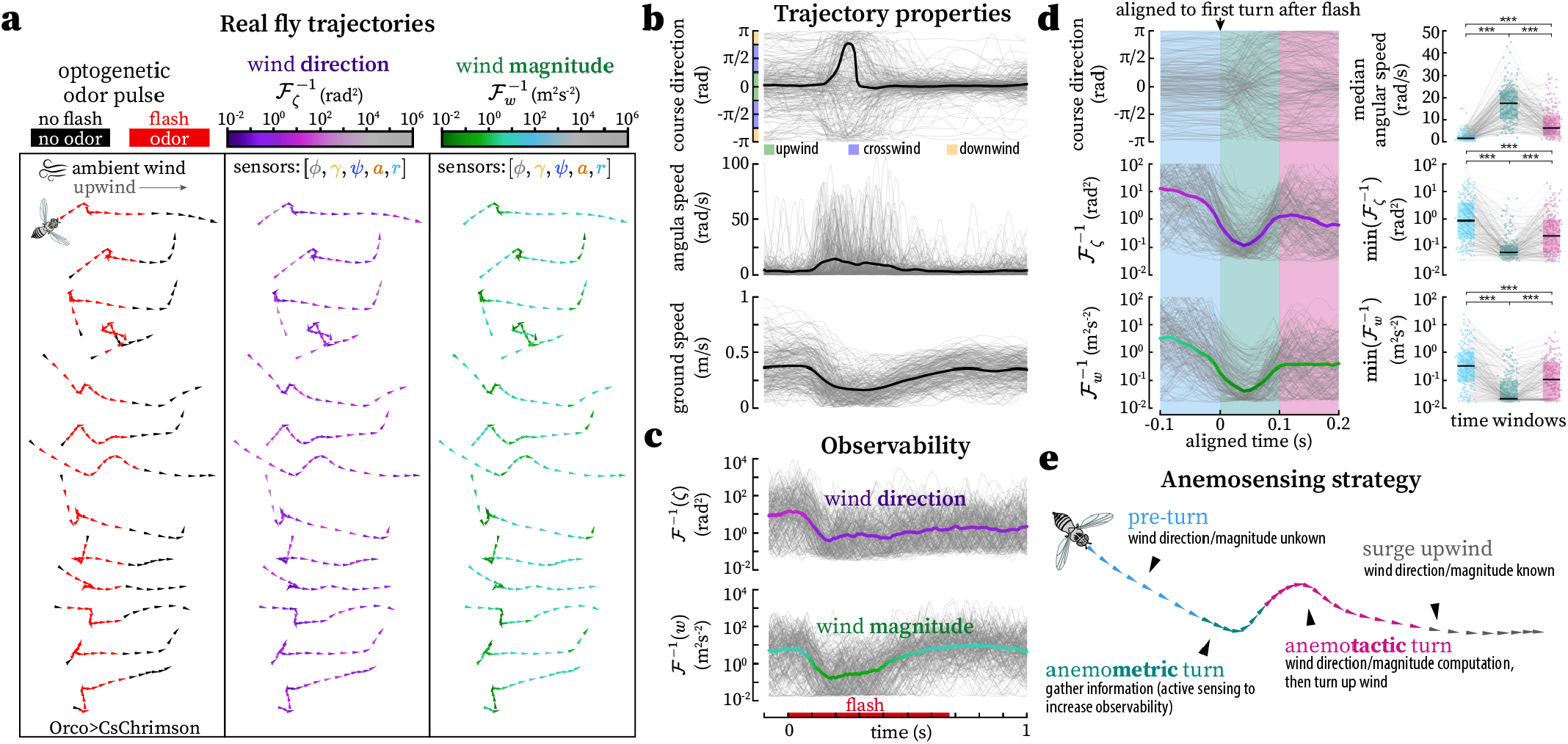
Observability properties of anemosensing behavior in measured fly flight. **a**. A subset of 2D trajectories of freely flying flies (Orco *>* CsCrimson) in a wind tunnel from Stupski and van Breugel ^32^ (left). A red light-sensitive ion channel in the antennae was optogenetically activated with a flash of red light for 0.675 s, which stimulated odor receptors and elicited anemosensing (see Methods). Each trajectory is 1.1 s long. The observability of the wind direction (middle, purples) and magnitude (right, greens) is shown assuming flies have access to heading, optic flow angle and magnitude, and apparent airflow angle and magnitude (see Figure 2 for variable legend). **b**. The global course direction, angular speed, and ground speed over time for *n* = 219 trajectories (gray). Black indicates the mean (or circular mean for course direction). **c**. The corresponding minimum error variance of wind direction and magnitude for each trajectory, assuming a 40 ms time window corresponding to 4 discrete measurements (*ω* = 4). Thick green and purple lines are the median. **d**. The course direction and minimum error variance aligned to the first turn detected after the optogenetic flash stimulus. Box plots illustrate the distribution of median angular speeds and minimum error variance before the turn (light blue) during the turn (green), and after the turn (pink). * * * *p <* 0.001 for t-test with Bonferroni correction (log transform applied for minimum error covariance data). *n* = 204 trajectories with a detectable first turn. **e**. Illustration of a proposed anemosensing strategy taking advantage of the increase in observability during a turn before orienting upwind.

To explicitly demonstrate that the first turn is well suited for estimating wind properties, we applied our observability analysis to this data. We assumed flies have access to heading *ϕ*, apparent airflow angle *γ* and magnitude *a*, ground velocity angle *ψ*, and optic flow magnitude *r*. We found that the first turn, which was also accompanied by a deceleration, was associated with a large increase in the observability of wind direction that would likely be sufficient to make an accurate wind direction estimate (Figure 5a–Figure 5c). Thus, flies exhibited characteristics of both the turning and acceleration/deceleration motif (Figure 2), one of the most observable motif combinations (Figure 4). By aligning the trajectories to the onset of the first turn we discovered that nearly all trajectories (93% that had a detectable first turn) displayed increased observability in the 100 ms following the turn onset, making this the most favorable time for a fly to estimate wind direction (Figure 5d). Furthermore, 70% of first turns crossed through either up- or down-wind and those that did not had a median minimum distance to upwind/downwind of 15^°^, which is also beneficial for wind direction observability (Figure 2e). The wind magnitude observability followed a similar increasing trend to the wind direction observability, supporting the claim that flies may be able to use wind magnitude information to to modulate their olfactory search ^32^. Our findings provide a computational basis for the use of an active sensing turn/deceleration (anemometric) before an upwind turn (anemotactic) during active anemosensing (Figure 5e).

### Exploiting active sensing motifs to filter sporadic state estimates from an artificial neural network

Although it would be possible for organisms and machines to move in a manner that would continuously ensure observability of states of interest, this would be an energetically inefficient approach and might be incompatible with some behavioral tasks. Data across taxa suggests that animals alternate between exploitation and exploration phases to gather information only when it is needed ^44^. How, then, might animals (or machines) ensure that they have continuous access to estimates of mission critical states? Observability information can be incorporated into a conventional Kalman filter ^17^ to improve estimation accuracy ^45^. The Kalman framework requires an accurate and dynamical model, can be computationally burdensome for high dimensional systems, is best suited for systems with full state observability, and does not exhibit direct analogs to how nervous systems process information. There is, however, evidence that animals will modulate sensory processing based on states such as angular velocity ^46^. With this as inspiration, we introduce a “motif informed filter” (MIF) framework that exploits knowledge of which motifs are correlated with observability to filter a single state estimate from an artificial neural network.

The general form of the MIF update equation is

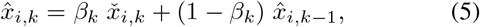

where 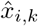 is the current filtered estimate of the state *x*_*i*_, 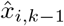 is the filtered estimate at the previous time-step, and 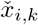 is the current raw estimate of the state. The raw estimate 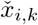 comes from an estimator function ℋ_*i*_ that depends only on the current and past measurements (and, optionally, inputs) from a time window of length *ω*,

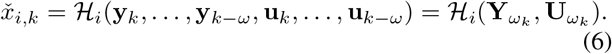

The coefficient *β*_*k*_ is a time-varying weight between 0 1 that determines how much to update the current filtered estimate with the current raw estimate. If observability can be calculated online, *β*_*k*_ can be determined from a scaled function of the observability of *x*_*i,k*_ (for statistical consistency, *β*_*k*_ should be equal to the constructability instead), for example:

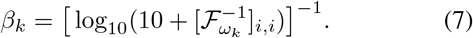

In practice, calculating observability online may be impractical, or impossible. For a more biologically plausible implementation *β*_*k*_ can be assigned a value based on the agents knowledge of the motifs exhibited in the window *ω*_*k*_ and their correlation to the observability of *x*_*i,k*_,

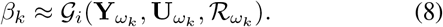

𝒢_*i*_ should be defined such that the value of *β*_*k*_ is near 1 when *ω*_*k*_ contains motifs correlated with high observability (i.e., *x*_*i,k*_ is highly observable) and near 0 in the absence of any observability correlated motifs (i.e., *x*_*i,k*_ is unobservable). We include the block diagonal sensor covariance matrix ℛ to account for the influence of sensor noise on the Fisher information: if the measurements are noisy it may be beneficial to update slowly, even during motifs corresponding to periods of high observability. The tools for discovering active sensing motifs from the prior sections can help inform the design of the function 𝒢. For instance, if the observability of *x*_*i*_ is strongly correlated with a rapid change in heading and a window size of 1 is sufficient for high observability, then the simple linear mapping *β*_*k*_ = *σ y*_*j,k*_ would be appropriate, where the change in heading 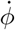 is encoded by measurement *y*_*j*_, and *σ* is a scaling factor that normalizes *β* and accounts for the sensor noise. However, other motifs may require more complex nonlinear functions.

To demonstrate our proposed MIF framework, we applied it to our anemosensing insect model and data. First, we designed a raw estimator (see Eq. 6) to estimate wind direction 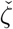 given noisy sensory measurements **y** = [*ϕ, γ, ψ*] (Figure 6a– Figure 6b). Although there are multiple approaches for estimating ζ from these measurements ^41^, we chose to use a feed-forward artificial neural network (ANN) for its generality, flexibility, and ability to operate without relying on a predefined dynamical model (Figure 6c). We created training data for our ANN by randomly simulating a diversity of trajectories and curating the ANN inputs (**y**) and output (*ζ*) from these trajectories (see Methods). We validated that the ANN could reasonably predict wind direction for withheld testing trajectories (Figure 6f, Figure S2). However, an important point is that we did not excessively try to optimize our ANN accuracy by adjusting hyperparameters, such as layer size or batch size, because we go on to show that leveraging observability can make up for a general lack of accuracy and optimization.

**Figure 6.**
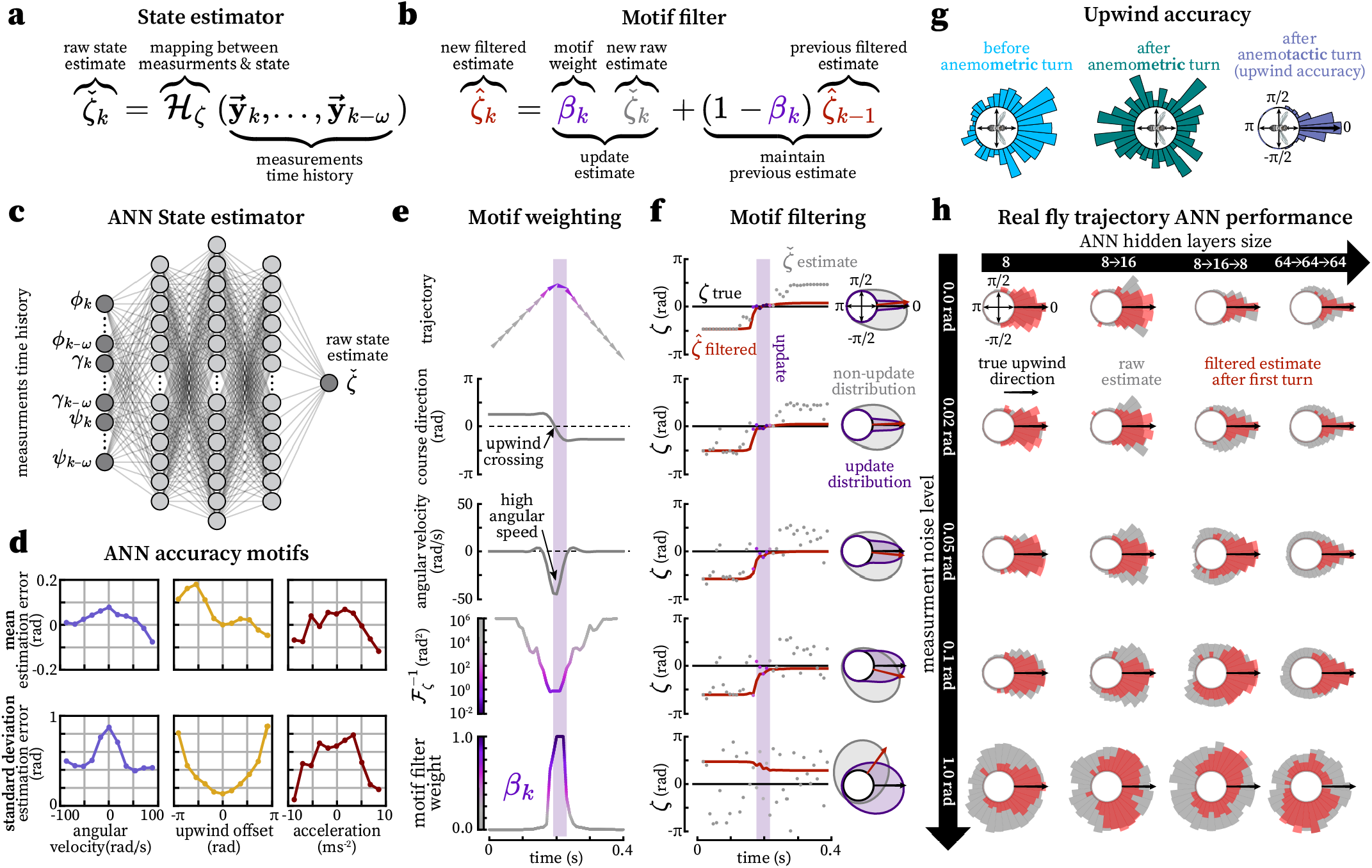
Motif informed filtering of artificial neural network estimates leverages observability to improve accuracy and persistence. **a**. The general form of an estimator that uses a time history of measurements **Y**^*ω*^ in the window from *k* to *k* − *ω* to estimate the current state, in this case the wind direction *ζ*. **b**. The general form of a motif informed filter (MIF) that weights incoming raw state estimates (from the estimator in *a*) with the previous filtered estimate. The time-varying variable *β*_*k*_ encodes the motif strength (analogous to observability), and thus how much to update the new filtered estimate. **c**. Illustration of a feed-forward artificial neural network (ANN) for wind direction estimation, in the form from *a*. The inputs are a time history of heading *ϕ*, apparent airflow angle *γ*, and ground velocity angle *ψ*. The current wind direction ζ is the only output. See Methods for specifics on network design and structure. **d**. The wind direction estimation error (true - predicted) mean and standard deviation as a function of trajectory properties for an ANN estimate with three 64 neuron layers and measurement noise with a standard deviation of 0.1 rad (see Methods). **e**. An example simulated trajectory with a turn passing through upwind that increases the observability (see Figure 2). The motif weight *β* (bottom) inversely scales with the observability. **f**. The true (*ζ*, black), ANN estimate (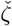, grey), and motif filtered estimate (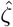, red) of wind direction for increasing levels of measurement noise. The ANN estimate was for a network with three 64 neuron layers (see Methods). The purple shaded region indicates where the motif weight *β* was above 0.7, corresponding to a 70% update in the MIF. The circular distributions on the right represent the corresponding raw estimate during high motif weight (*β >* 0.7, purple) and during low motif weight (*β <* 0.7, grey). The black arrow points to the the true ζ and the red arrow points towards the motif filtered estimate after the turn. **g**. The distribution of real fly trajectory headings (from Figure 5) before the anemometric turn (light blue), after the anemometric turn (green), and after the anemotactic turn (teal). Note that the distribution after the anemotactic turn is largely pointed towards the true upwind direction (black arrow), indicating that flies are capable of successfully estimating and navigating upwind. (*n* = 219 trajectories). **h**. Results of the MIF applied to real fly trajectories. The circular distributions of the raw estimate from the ANN (grey) compared to the motif filtered estimate (red) after the first spike in *β* (exceeded and then dropped below 0.4), corresponding to a turn. This analysis was repeated for four different ANN’s (trained on identical data) and with different levels of sensor noise (the ANN’s were all trained with measurement noise with a standard deviation of 0.01 rad).

First, we confirmed that our ANN accuracy was correlated with previously discovered motifs. Specifically, larger changes in course direction (i.e. turning), smaller upwind offsets (computed from measurements as *ψ* − *γ*), and larger accelerations and decelerations were all found to decrease the estimation error 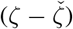 mean and standard deviation (Figure 6d). This is an important verification step because an ANN estimator is not necessarily ideal and thus is not guaranteed to exploit all observable motifs.

Next, we simulated new trajectories with fly-like rapid turns, applied varying levels of noise to the measurements, and used our ANN to estimate 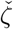 (Figure 6e). These raw wind direction estimates had large variances and errors (Figure 6f). This is unsurprising since the network was not trained these specific trajectories with this type of turning motif, we used higher levels of noise compared to our training set, and the trajectories had many regions of low observability. However, the mean and variance were dependent on the trajectory itself and there were sparse regions where 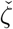 was accurate and precise. These regions corresponded closely with the increased observability during the motifs we discovered (Figure 6f, Figure 2). Specifically, instances where a fly’s heading crossed through upwind (0^°^) at sufficient angular speed led to the most accurate and least variable estimates (Figure 6f), as predicted by our prior observability analysis (Figure 2, Figure 4). To maintain a persistently accurate state estimate that is not corrupted by the periods of low observability we now apply the MIF framework.

To use the MIF, we designed a function 𝒢 to assign a time-varying value to *β*_*k*_ (Figure 6b) that incorporated the turning and upwind offset motifs (see Methods for specific details). We did not use the acceleration motif for this example because acceleration is not available from the limited sensor group we highlight in this example, but it could be included either as another measurement in the system or through a control input.

We found that this motif weight function was sufficient to capture the observability of wind direction associated with simulated turns and was thus correlated with the regions of accurate and precise estimates from the ANN (Figure 6e). We then applied the MIF (Figure 6b) which led to the filtered estimate 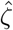 smoothly converging to the true of wind direction ζ during the turn, and maintaining this estimate thereafter (Figure 6f). Notably, the MIF performance estimate was accurate and precise until relatively large (*>* 0.1 rad) noise levels, which is expected as noise levels bound the observability minimum estimation error covariance (Eq. 2).

As a final test, we applied the MIF framework to real fly trajectories (see Figure 5). Flies are able to quickly orient upwind after a brief odor stimulus with impressive accuracy, implying that they compute a similarly accurate estimate of the wind direction (Figure 6g). Therefore, we wanted to see if it was possible to match their behavioral accuracy with an ANN and MIF. We replayed the simulated measurements corresponding to all of our real fly trajectories into ANN’s with different hidden layer structures (but trained on the same simulated dataset) and added varying levels of measurement noise (Figure 6h). We found that after applying our MIF ANN’s with larger and/or more hidden layers yielded distributions of 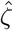 that matched or exceeded the distributions of fly course directions, even for relatively high noise levels (Figure 6h, Figure 6g). However, without the MIF, the ANN always under performed the real flies. Although a larger ANN with access to more time history might be able to match our MIF performance, our MIF framework offers an approach for maximizing the performance of small networks. In fact, our ANN with three small layers and 32 total neurons (8 + 16 + 8 = 32) matched the fly performance at low-to-medium noise levels, suggesting that real insects may not require a large wind direction estimation circuit if they incorporate an approach similar to our motif informed filter.

## DISCUSSION

We introduced a novel framework we call “BOUNDS: Bounding Observability for Uncertain Nonlinear Dynamic Systems” for empirically evaluating the observability of individual state variables that has particular advantages over prior methods. By considering the noise levels associated with sensors, our approach is able to offer a physical unit for observability that describes a bound on how well an ideal estimator could estimate a given state variable. Our approach is amenable to high dimensional and/or partially observable nonlinear systems. Furthermore, our method can analyze real world trajectory data in conjunction with a model, which is useful for evaluating data collected from moving organisms or machines. We showed how our framework can be applied to manually or methodically discover active sensing motifs for a model system (anemosensing). For higher order systems it may be beneficial to take a more automated approach for discovering motifs using techniques like genetic algorithms, particle swarm optimization, or reinforcement learning. Lastly, we presented a framework for integrating observability, or correlated motifs, to intelligently filter state estimates. Altogether, our approach provides a means to model active sensing and estimation in organisms and machines.

### Tips and Tricks

Our approach has just two primary hyper-parameters, 1) the regularization parameter *λ* (Eq. 4), and 2) the window size *ω* over which to analyze the observability. It is best practice to choose *λ* to be large enough to numerically invert the Fisher information matrix, but small enough to not distort the total information. In general, a good guideline is to choose *λ* to be less than the smallest nonzero eigenvalue of the Fisher information matrix ^47^. Alternatively, one could symbolically compute the limit in Eq. 4, however this can be computationally expensive and can yield infinite values corresponding to unobservable states. For *ω* we recommend choosing a window size roughly equal to the fastest behavioral motif of interest. Other practical suggestions for applying our approach are provided in the Methods.

### Relation to constructability and inclusion of model uncertainty

We chose to analyze observability, rather than the dual concept of constructability (which analyzes the final state, rather than the initial state ^28^) because it allowed us to use an initial state perturbation approach for empirically calculating the observability matrix 𝒪. This perturbation approach is efficient, and works seamlessly with non-closed form models. For the purpose of discovering active sensing motifs, both approaches will provide equivalent insight. In cases where a quantification of the error bounds for the final state are necessary, our calculations of ℱ can be replaced with an iterative equation that uses a discretized time-varying approximation of the continuous time dynamics ^28^. Taking this approach would also allow for the inclusion of non-zero process noise matrix *Q* to understand how uncertainty of an agent’s internal model affects the observability (or constructability) of states of interest.

### Exploiting observability for estimation

Perhaps the most critical component to active sensing it knowing *when* to do it. Constantly performing active sensing maneuvers can be wasteful, or may conflict with other navigation goals. To achieve a balance many animals appear to alternate between these two goals in an exploration vs exploitation tradeoff that is triggered when estimates reach a threshold of error ^44^. The motif informed filter we introduce provides a flexible and parsimonious framework for taking advantage of observability (or an approximation thereof) information to intelligently update state estimates only during periods of intermittent active sensing (Figure 6). In our example we used a feed-forward ANN for the raw estimator ℋ, and empirically designed a motif weighting function 𝒢, however with sufficient training data both functions could be performed by two separate ANNs. We took this approach to illustrate how the MIF framework can be used to provide accurate estimates of an individual state variable. This approach could be extended to full state estimates, and to include the effects of known dynamics between active sensing bouts, by using a partial-update Schmidt Kalman filter ^45^ that adjusts the Kalman gain proportionally to the level of observability for each state. Alternatively, a recurrent neural network (RNN) that can simultaneously estimate and filter might able to perform both steps at once without the need for an explicit model ^48,49,50,51^. Our observability framework might help with training an RNN by providing scaffolded learning objectives to help the RNN discover useful motifs more quickly.

### Insights into anemosensing and processing in the fly brain

We applied our observability tools to discover candidate active sensing motifs for anemosensing in flying insects (Figure 2– Figure 4). In general, we found that large turns and decelerations enrich information about wind direction and magnitude. We also show that in dynamic wind both passive changes in course and gust rejection strategies can provide sufficient information for estimation of the wind vector. Although the quantitative values of our calculations depend on the accuracy of the model we used, we expect the general principles will remain valid when considering a more detailed 3D biomechanically accurate model. Supporting our hypotheses, we show that flies employ the turning and deceleration motifs at times where estimating wind direction is of high priority (Figure 5). Determining whether flies do indeed estimate wind direction during these maneuvers, and discovering which sensor combinations they use, will require targeted silencing and imaging experiments. Our results show that there are many partially overlapping groups of sensors can be used to estimate wind direction, raising the possibility that insects could potentially flexibly remap or reweight how cues are integrated in a context dependent manner.

During both the upwind surge and casting states of olfactory navigation flies would benefit from maintaining a persistent wind direction estimate that is only updated during time windows when it is observable. Do flies use a strategy similar to our motif informed filter to achieve this? Though we cannot be certain, the concept is feasible. Dopaminergic neurons innervating the head direction network in the central complex are selectively active during changes in heading, and modulate processing (in this case, they promote plasticity) ^46^. We suggest that a similar mechanism might be used elsewhere in the brain to modulate when persistent estimates are updated, effectively using a neuromodulator to encode an anlagous *β*_*k*_ parameter in our motif informed filter. Over time, the fly brain may have evolved to recognize that there is more information available about certain states during specific maneuvers, and modulate the filtering of those state estimates during those periods.

## Conclusion

We suggest that many of the movements animals make may serve as opportunities for information gathering, and machines could exploit similar movements with proper estimator design. In most cases, active sensing motifs are likely to be turning or changes in ground speed, either of the body, or the sensors themselves. Such maneuvers could help to make up for a lack of sensors or sensor quality, potentially reducing the metabolic or fabrication costs. In our example we show that such maneuvers can aid in estimation of ambient wind direction, but similar movements would help with estimation of other external forces, and even for estimating parameters of the agents body such as its mass, drag coefficients, muscle force coefficients, and sensor calibration coefficients. Our approach provides a rigorous computational framework for exploring this hypothesis. The methods we describe here can also be used to provide insight into how they might implement an estimator. In this paper we considered sensors that report pure measurements of individual states, or simple combinations (e.g. optic flow). However, brains often combine different sensory signals together, often with low- or high-pass filter dynamics. Our approach can be used to understand which state variables these multiplexed signals can estimate. Alternatively, it is possible to use our methods to discover how sensory signals can be multiplexed so as to minimize the number of channels needed to convey the information necessary to estimate quantities of interest. In contrast to other sensor compression approaches our method will allow for higher levels of compression in exchange for requiring the use of an active sensing motif. Collectively, our framework will help decode active sensing at the behavioral and neural level in organisms, and inspire the design of intelligent active sensing in machines.

## METHODS

### Fly-wind dynamical model

#### Model

To investigate active sensing in in the context of anemosensing we used a continuous-time model of a flying insect’s flight dynamics based on previous work ^38^

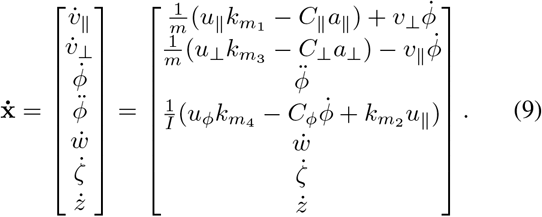

Although we used a continuous-time model to describe the underlying dynamics, the dynamics were effectively converted to discrete-time when solving the differential equations (see next section). This model accounts for a flying insect agent’s translation and rotation in a 2D plane in the presence of ambient wind. The state variables include the ground velocity components parallel *v*_‖_ and perpendicular *v*_⊥_ to the agents heading orientation, heading *ϕ*, angular velocity 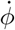, ambient wind magnitude *w* and direction *ζ*, and elevation *z*. The agent controls translation by flight forces through the center of mass and the heading is controlled by a turning torque. The corresponding controls for each degree of freedom are **u** = [*u*_‖_, *u*_⊥_, *u*_*ϕ*_], which allow any arbitrary trajectory to be simulated with proper control, as in Figure 1a. Our model also accounts for dynamics from mass *m* and inertia *I*, and airflow induced translational *C*_‖,⊥_ and rotational drag *C*_*ϕ*_ that arises from a combination of motion induced airflow and ambient wind. Parameter values were chosen based on those measured or estimated for *Drosophila* (Table 1). For observability calculations, we included the mass and drag parameters in Table 1—as well as motor calibration parameters 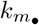 —as auxiliary state variables ^38^. This allowed us to specifically study what happens when an insect does not have a perfect model of its flight dynamics. All motor calibration parameters were set to 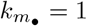 except 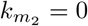.

**Table 1.** *Drosophila* parameters. Note that the translation damping was computed from the time constant (*τ* = 0.170s) ^54^ and the mass as *C*_‖,⊥_ = *m/τ*.

| Parameter | Value | Source |
| --- | --- | --- |
| mass ($m$) | $2.5 \times 10^{-7}$ kg | 52 |
| yaw inertia ($I$) | $5.2 \times 10^{-13}$ kg m<sup>2</sup> | 53 |
| translation damping ($C_{\parallel,\perp}$) | $1.5 \times 10^{-6}$ kg s<sup>-1</sup> | 54 |
| rotational damping ($C_{\phi}$) | $2.7 \times 10^{-11}$ Nms/rad | 53 |

The measurement dynamics of our model were chosen depending on which set of sensory cues we were investigating (Figure 2). However, all measurements were defined as a function of the states, and chosen from the set

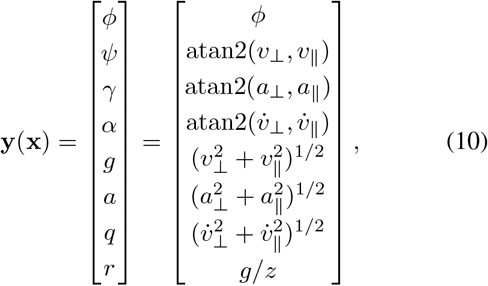

where the apparent airflow parallel and perpendicular components are

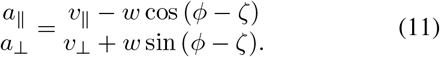

#### Simulation and control

We simulated our fly-wind dynamical model in Python with a nonlinear differential equation solver ^55^. Even though our model was defined with continuous time dynamics, we simulated each time-step discretely with constant inputs for each step. This was done to ensure that we could start from any given point along a trajectory and maintain the exact simulation thereafter, which was necessary to construct observability matrix in sliding windows.

We used model predictive control (MPC) with full state feedback to precisely control flight trajectories through our model (Eq. 9) (Figure 1b). In principle, other control methods could work, but MPC is a good general choice for controlling non-linear systems. We first prescribed—or took from measured flight trajectories—a desired set-point for *v*_‖_, *v* _⊥_, and *ϕ* over time (Figure 1a), then used a open-source MPC toolbox ^56^ to find the optimal control inputs to track the set-point variables. Our MPC cost function penalized the squared error between each of our set-point series while adding an additional penalty for each control input to ensure more naturalistic force and torque inputs. Specifics can be found in our publicly available code (CITE GITHUB).

### Observability calculations

#### Empirical observability matrix

We constructed empirical observability matrices 𝒪 as previously described ^15^. In brief, for a given state trajectory **x**_*k*_ we set a window size *ω* and initial state **x**_*k*_ in sliding windows (Figure 1a). Then we perturbed each individual initial state variable by a small amount *ε* = 1 × 10^−5^ in both directions 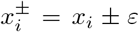. The resulting differences in the sensor measurements (Figure 1c) was collected and normalized by the perturbation amount Δ**y**_*k*_*/*2*ε* and arranged as rows in 𝒪 (Figure 1e). All angular measurement variables were unwrapped to ensure that Δ**y**_*k*_ did not have any discontinuities, a perquisite for empirical observability analysis. We added a small amount (0.01 m s^−2^) of translational acceleration to all simulated trajectories to ensure that the acceleration angle *α* was always defined—otherwise *α* would be discontinuous. We set a window size of *ω* = 4 for all simulations.

#### Fisher information and inverse

For each 𝒪 we computed the Fisher information matrix as ℱ = 𝒪^⊺^ℛ^−1^𝒪. We set the block diagonal noise covariance matrix ℛ= 0.1 *I*_(*p×ω*)×(*p×ω*)_ where *p* is the number of measurements and *ω* is the sliding window size. This assumes the noise variance on each measurement is either 0.1 rad for angular sensors or 0.1 m s^−1^ for magnitude sensors. The true noise properties of sensory measurements in *Drosophila* are currently unclear, however these values are reasonable guesses and do not have a large impact on our results because larger or smaller values ℛ of simply scale ℱ. We do not consider the process noise *Q* in this manuscript, but with some modifications to our method (see ^28^) this could be taken into account as well.

Calculating the inverse of ℱ is not straightforward due to the presence of unobservable directions in the fly-wind model, which results in a matrix that is not invertible. A pseudo inverse can be computed, but this results in misleading (often small, when they should be very large) values within the diagonal elements of ℱ^−1^ for unobservable states. Therefore, we chose to add a small amount of observability to each direction in ℱ to ensure computing a traditional inverse was possible (Eq. 4). We refer to this inverse as the “Chernoff inverse” ^31^, following the naming convention of prior literature on matrix inverses ^57^. Note that the regularization does not have to be of the form *λI*_*n×n*_. A more general regularization of *λJ*, where *J* is any *n × n* positive definite matrix, is allowable. For simplicity, we chose *J* to be the identity matrix. This method results in a cap on the upper end of the error noise covariance matrix equal to 1*/λ*. For all systems, *λI*_*n×n*_ must be carefully selected to ensure that it is small enough not to distort the true observability properties, while still large enough to avoid numerical instabilities associated with computing a numerical inverse. We set *λ* = 1 × 10^−6^, which resulted in error variance values above the relevant values computed for the fly-wind model. We normalized and inverted the error variance data for display in the radar plots in Figure 2–Figure 3 and tree graph Figure 5 by taking 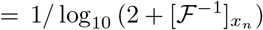, where 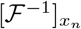 is the *n*^*th*^ diagonal element of ℱ^−1^, corresponding to the minimum error covariance for state *x*_*n*_.

#### Tips and tricks

In general, our method is appropriate for any system, that is as long as the system can be simulated. However, it is critical to ensure that none of the state variables have discontinuities in the state space of the simulation. Otherwise, artifacts can manifest as extremely large values in the observability matrix, which corrupt the computation of the Fisher information matrix. For instance, when dealing with polar variables and angles it is crucial to ensure that the state perturbations required to construct the observability matrix do not lead to angle wrapping discontinuities. Magnitude variables must not be perturbed to be negative, as magnitudes are defined to always be positive, because this would have the same effect in the observability matrix as applying a 180^°^ perturbation to the associated angular quantity. It is also important to ensure that all measurement functions are always defined. If a measurement consists of a fraction such as *y*_2_*/y*_2_, then *y*_2_ = 0 would lead to an undefined measurement which breaks observability assumptions.

When considering the observability associated with many different combinations of sensors, it is best practice to compute the observability matrix once for all the potential states and sensors, and for the largest time window to be considered. Then one can pull out smaller observability matrices corresponding to whatever combinations of states, sensors, and time windows for further analysis (Fisher information, etc.). This is in contrast to performing many simulations to reconstruct a new observability matrix for each unique combination, which effectively contains the same information. This approach avoids cumbersome computations and makes iterative investigation of active sensing motifs much faster, although this is dependent on the speed of the simulation required to construct the observability matrix.

### Fly data and processing

All measured *Drosophila* trajectories were procured from Stupski and van Breugel ^32^. The ground speed and course direction of each individual trajectory were used for the set-points for our MPC routine. Since heading data was not available in the published data, we assumed that the fly heading was aligned with their course direction at all times. We then simulated the trajectory with MPC and collected the corresponding sensory measurements. These simulations and measurements were used to perform observability analysis (Figure 5).

### Artificial neural network

We designed feed-forward artificial neural networks (ANNs) for regression, with multiple inputs corresponding to putative sensory measurements a fly might have access to. For Figure 6, the inputs were heading *ϕ*, apparent airflow angle *γ*, and ground velocity angle *ψ*. We augmented each input with a time-history *ω* of the measurements, where we set *ω* = 4 to be consistent with the observability analysis (Figure 6b). The structure of all the ANNs consisted of the input layer, one or multiple hidden layers, and an output layer with a single neuron for wind direction *ζ*. All neurons were fully connected and used a rectified linear unit activation function, except for the output layer which used a linear activation function. We built and trained our network using TensorFlow ^58^ and Keras ^59^. We designed a custom loss function to deal with the circular (discontinuous, and therefore harder to train) nature of the output variable:

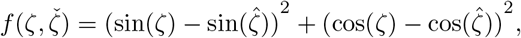

where ζ is the true wind direction of the training data and 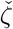 is the ANN output (predicted *ζ*). This function effectively penalizes the squared error between the sine and cosine components of ζ and 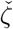.

We simulated a 11,000 trajectory dataset (Figure 6a) to train and test the ANNs. We randomly chose the simulated trajectory and wind properties from a uniform distribution between a set range of values

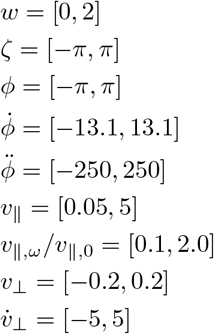

where *v*_‖,*ω*_*/v*_‖,0_ represents the ratio of initial and final *v*_‖_ (Figure S2a). This was used instead of 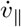 to ensure flight was always generally directed forward and there was not backwards flight for small starting *v*_‖_ values. We applied a 80-20 train-test split to the data. Each trajectory was 0.2 s (21 discrete points). We augmented the measurements from each trajectory with *ω* = 4 prior time-steps, so only the 17 points with *ω* prior points available were used in training/testing. We trained each ANN for 1000 epochs with a batch size of 256, which was sufficient for the validation loss to converge (Figure S2b– Figure S2d). We also added a Gaussian noise layer with a mean of 0 and standard deviation of 0.01 rad after the input layer, which was only active during training, to help the ANN deal with noise in measurements.

While many hidden layer structures are sufficient for wind direction estimation, we considered a few different structures to highlight the tradeoffs of a more or less accurate network when combined with motif informed filtering (Figure 6h, Figure S2). We started with a simple 8 neuron hidden layer, which was not sufficient to achieve an accurate estimate, even during observable regions (Figure 6h). Adding a second 16 neuron hidden layer only modestly improved the results (Figure 6h). However adding a third layer of 8 neurons (total hidden layers of 8, 16, 8 neurons) allowed an accurate estimate to be made in the observable regions for lower levels of measurement noise (Figure 6h). The largest network we tested consisted of three 64 neuron hidden layers, which yielded the most accurate and precise estimates during observable regions, and was the most robust to measurement noise with motif filtering (Figure 6h).

### Motif informed filter

We prescribed an exponential scaling for the turning motif weight *β*_*k*,turn_, wherein large turns led to values near 1 and small turns led to *β*_*k*_ near 0

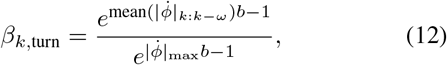

where *b* was the strength of the exponential. We defined a binary function for the upwind offset motif weight *β*_*k*,upwind_, where *β*_*k*,upwind_ = 1 if there was an upwind crossing within the time window, which we detected from the measurements by checking for *ψ* − *γ* to change sign, and *β*_*k*,upwind_ = 0 if there was not an upwind crossing. We then fused these functions to set the total MIF weight as

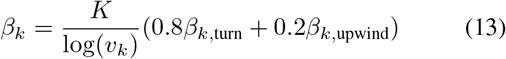

where 80% of the weight was controlled by the turning motif and 20% of the weight was controlled by the upwind crossing motif. The total weight was inversely scaled with the noise level *K/*log(*v*_*k*_), where *v*_*k*_ is the noise standard deviation in rad and *K* is a scaling factor that mapped the normalized the weight to be 0 when *v*_*k*_ ≥ 0.5 rad and 1 when *v*_*k*_ ≤ 0.01 rad.

## DATA AND CODE AVAILABILITY

We provide a python package to implement BOUNDS (https://github.com/vanbreugel-lab/pybounds) which will be updated periodically. Specific data associated with this manuscript will be made public upon publication.

## FUNDING

This work was supported by a National Science Postdoctoral Fellowship in Biology to Benjamin Cellini (NSF 22-623). The work was also supported by the Air Force Office of Scientific Research (FA9550-21-0122) to FvB, the NSF AI institute in dynamics (2112085) to FvB, NSF-EFRI-BRAID (2318081) to FvB, Sloan Research Fellowship (FG-2020-13422) to FvB, and DEPSCoR (FA9950-23-1-0483) to FvB. For the purpose of open access, the author has applied a CC BY public copyright license to any Author Accepted Manuscript version arising from this submission.

## CONFLICTS OF INTEREST

The authors declare no conflict of interest.

## AUTHOR CONTRIBUTIONS

Conceptualization: B.C, B.B, F.v.B.; Methodology: B.C, B.B, F.v.B.; Software: B.C; Formal analysis: B.C, S.D.S; Investigation: B.C, S.D.S, F.v.B.; Resources: F.v.B.; Data curation: S.D.S; Writing - original draft: B.C, F.v.B.; Writing - review & editing: B.C, B.B, S.D.S; F.v.B.; Visualization: B.C, F.v.B.; Supervision: F.v.B.; Project administration: F.v.B.; Funding acquisition: B.C, F.v.B.;

## SUPPLEMENTARY MATERIAL

**Supplementary Figure S1.**
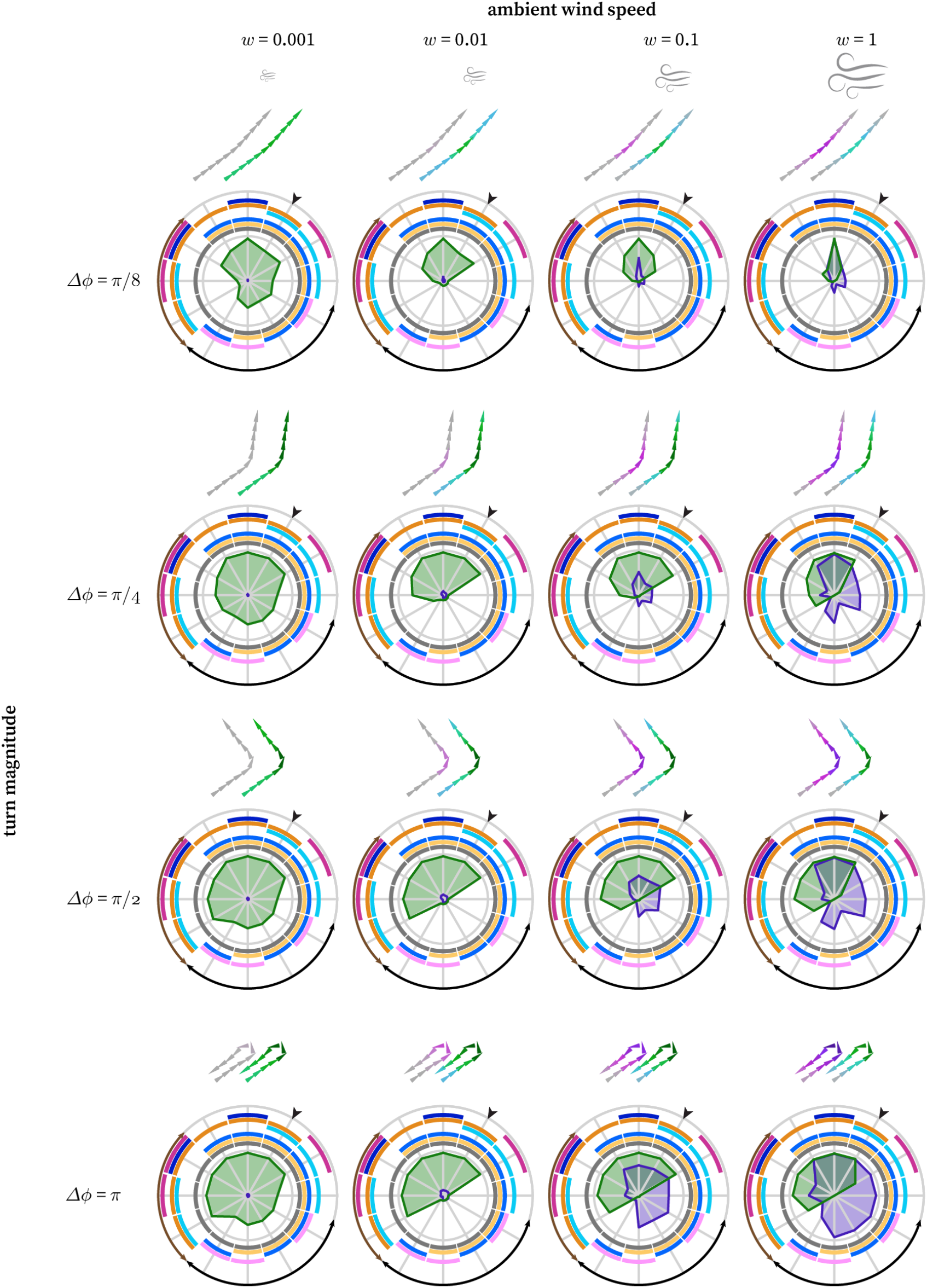
Interactions between wind speed and tuning magnitude. The effects of wind speed and turn magnitude on wind direction and magnitude observability for various sensor groupings. The ground speed was kept constant at 0.3 m s^−1^. The wind speed was kept constant within each simulation, but varied across simulations (no dynamic wind). See Figure 2 for legend and variable definitions.

**Supplementary Figure S2.**
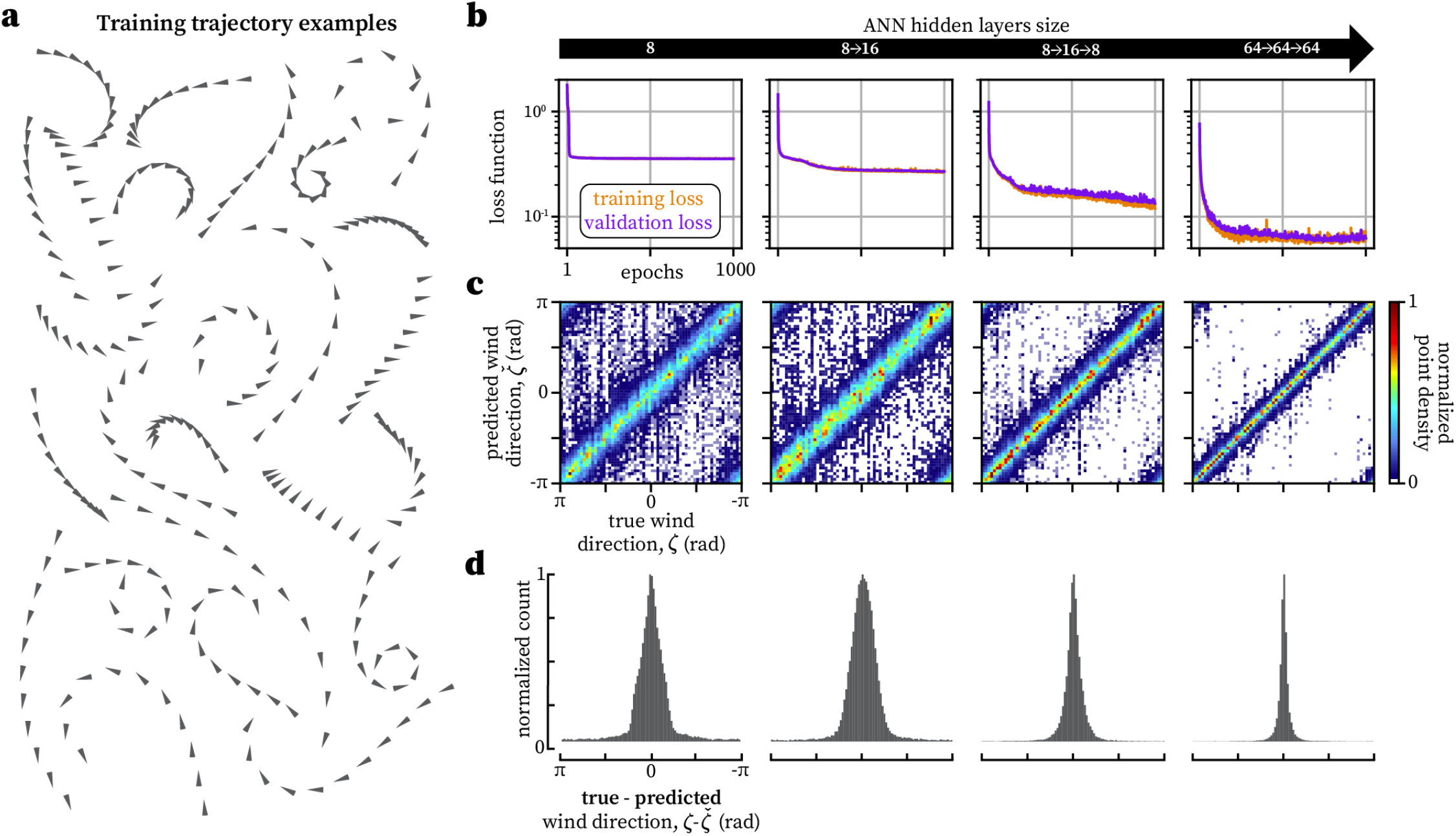
ANN training and accuracy. **a**. A small subset of example simulated training trajectories used to train artificial neural networks (ANN) for wind direction estimation. **b**. The training (orange) and validation (purple) loss over 1000 epochs. A 80–20 validation split was used. **c**. True wind direction (*ζ*) vs predicted wind direction 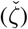 for ANN’s with different hidden layer structures. **d**. Distribution of errors 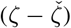 for each ANN.

## Notes

### Competing Interest Statement

The authors have declared no competing interest.

## REFERENCES

[1] Ryan J. Rowekamp and Tatyana O. Sharpee. Cross-orientation suppression in visual area V2. Nature Communications, 8:1–9, 2017. doi:10.1038/ncomms15739.

[2] Shaoqing Ren, Kaiming He, Ross Girshick, and Jian Sun. Faster R-CNN: Towards Real-Time Object Detection with Region Proposal Networks. IEEE Transactions on Pattern Analysis and Machine Intelligence, 39(6):1137–1149, jun 2017. doi:10.1109/TPAMI.2016.2577031.

[3] Scott Cheng-Hsin Yang, Daniel M. Wolpert, and MátéLengyel. Theoretical perspectives on active sensing. Current Opinion in Behavioral Sciences, 11:100–108, oct 2016. doi:10.1016/j.cobeha.2016.06.009.

[4] Floris van Breugel, Kristi Morgansen, and Michael H. Dickinson. Monocular distance estimation from optic flow during active landing maneuvers. Bioinspiration & Biomimetics, 9(2):025002, may 2014. doi:10.1088/1748-3182/9/2/025002.

[5] Bryson Lingenfelter, Arunava Nag, and Floris van Breugel. Insect inspired vision-based velocity estimation through spatial pooling of optic flow during linear motion. Bioinspiration & Biomimetics, 2021. doi:10.1088/1748-3190/ac1f7b.

[6] Karl Kral. Side-to-side head movements to obtain motion depth cues:. Behavioural Processes, 43(1):71–77, apr 1998. doi:10.1016/S0376-6357(98)00007-2.

[7] Philip RL Parker, Elliott TT Abe, Natalie T Beatie, et al. Distance estimation from monocular cues in an ethological visuomotor task. eLife, 11:e74708, sep 2022. doi:10.7554/eLife.74708.

[8] Mitra J. MJ Mitra J. Hartmann. Active sensing capabilities of the rat whisker system. Autonomous Robots, 11 (3):249–254, 2001. doi:10.1023/A:1012439023425.

[9] Tanvi Deora, Mahad A Ahmed, Thomas L Daniel, and Bing W Brunton. Tactile active sensing in an insect plant pollinator. Journal of Experimental Biology, 224 (4):jeb239442, 2021.

[10] Benjamin Cellini, Wael Salem, and Jean-Michel Mongeau. Complementary feedback control enables effective gaze stabilization in animals. Proceedings of the National Academy of Sciences, 119(19):e2121660119, may 2022. doi:10.1073/pnas.2121660119.

[11] Benjamin Cellini and Jean-Michel Mongeau. Active vision shapes and coordinates flight motor responses in flies. Proceedings of the National Academy of Sciences, 117(37):23085–23095, sep 2020. doi:10.1073/pnas.1920846117.

[12] Reinhold Necker. Head-bobbing of walking birds. Journal of comparative physiology A, 193:1177–1183, 2007.

[13] Melville J Wohlgemuth, Ninad B Kothari, and Cynthia F Moss. Action enhances acoustic cues for 3-d target localization by echolocating bats. PLoS biology, 14(9): e1002544, 2016.

[14] Marie P. Suver, Ashley M. Medina, and Katherine I. Nagel. Active antennal movements in Drosophila can tune wind encoding. Current Biology, 33(4):780–789.e4, feb 2023. doi:10.1016/j.cub.2023.01.020.

[15] Benjamin Cellini, Burak Boyacioğlu, Floris van Breugel, et al. Empirical Individual State Observability. 2023 62nd IEEE Conference on Decision and Control (CDC), pages 8450–8456, ec 2023. doi:10.1109/CDC49753.2023.10383812.

[16] Abhinav Kunapareddy and Noah J. Cowan. Recovering Observability via Active Sensing. In 2018 Annual American Control Conference (ACC), volume 2018-June, pages 2821–2826. IEEE, jun 2018. doi:10.23919/ACC.2018.8431080.

[17] R.E. Kalman. On the general theory of control systems. IFAC Proceedings Volumes, 1(1):491–502, aug 1960. doi:10.1016/S1474-6670(17)70094-8.

[18] Sarah A. Stamper, Eatai Roth, Noah J. Cowan, and Eric S. Fortune. Active sensing via movement shapes spatiotemporal patterns of sensory feedback. Journal of Experimental Biology, 215(9):1567–1574, may 2012. doi:10.1242/jeb.068007.

[19] Debojyoti Biswas, Luke A. Arend, Sarah A. Stamper, et al. Closed-Loop Control of Active Sensing Movements Regulates Sensory Slip. Current Biology, 28(24):4029–4036.e4, dec 2018. doi:10.1016/j.cub.2018.11.002.

[20] Eduardo D Sontag, Debojyoti Biswas, and Noah J Cowan. An observability result related to active sensing. arXiv preprint, pages 1–12, oct 2022. doi:10.48550/arXiv.2210.03848.

[21] Sanjay Lall, Jerrold E Marsden, and Sonja Glavaški. Empirical model reduction of controlled nonlinear systems. IFAC Proceedings Volumes, 32(2):2598–2603, jul 1999. doi:10.1016/s1474-6670(17)56442-3.

[22] James O. Berger. Statistical Decision Theory and Bayesian Analysis. Springer Series in Statistics. Springer New York, New York, NY, 1985. doi:10.1007/978-1-4757-4286-2.

[23] Harald Cramér. Mathematical Methods of Statistics. Princeton Univ. Press, Princeton, NJ, 1946.

[24] Abhay K. Singh and Juergen Hahn. On the use of empirical gramians for controllability and observability analysis. Proceedings of the American Control Conference, 1 (May):140–141, 2005. doi:10.1109/acc.2005.1469922.

[25] Arthur J. Krener and Kayo Ide. Measures of unobservability. In Proceedings of the 48h IEEE Conference on Decision and Control (CDC) held jointly with 2009 28th Chinese Control Conference, pages 6401–6406. IEEE, ec 2009. doi:10.1109/CDC.2009.5400067.

[26] Roman Vaxenburg, Igor Siwanowicz, Josh Merel, et al. Realistic physics simulation of fruit fly locomotion with reinforcement learning. arXiv preprint, 2024.

[27] E. F. Camacho and C. Bordons. Model Predictive control. Advanced Textbooks in Control and Signal Processing. Springer London, London, 2007. doi:10.1007/978-0-85729-398-5.

[28] Burak Boyacioğlu and Floris van Breugel. Duality of Stochastic Observability and Constructability and Their Relation to the Fisher Information. 2024. doi:arXiv:2410.19975.

[29] Chun Sik Hwang. Observability and Information Structure of Nonlinear Systems,. PhD thesis, Oregon State University, 1985.

[30] Claude Jauffret. Observability and fisher information matrix in nonlinear regression. IEEE Transactions on Aerospace and Electronic Systems, 43(2):756–759, apr 2007. doi:10.1109/TAES.2007.4285368.

[31] Herman Chernoff. Locally optimal designs for estimating parameters. The Annals of Mathematical Statistics, pages 586–602, 1953.

[32] S David Stupski and Floris van Breugel. Wind gates olfaction-driven search states in free flight. Current Biology, pages 1–15, jul 2024. doi:10.1016/j.cub.2024.07.009.

[33] Marie P. Suver, Andrew M.M. Matheson, Sinekdha Sarkar, et al. Encoding of Wind Direction by Central Neurons in Drosophila. Neuron, 102(4):828–842.e7, may 2019. doi:10.1016/j.neuron.2019.03.012.

[34] Yvette E. Fisher. Flexible navigational computations in the Drosophila central complex. Current Opinion in Neurobiology, 73:102514, apr 2022. doi:10.1016/j.conb.2021.12.001.

[35] Cheng Lyu, L. F. Abbott, and Gaby Maimon. Building an allocentric travelling direction signal via vector computation. Nature, 601(7891):92–97, jan 2022. doi:10.1038/s41586-021-04067-0.

[36] Tatsuo S. Okubo, Paola Patella, Isabel D’Alessandro, and Rachel I. Wilson. A neural network for wind-guided compass navigation. Neuron, 107(5):924–940.e18, September 2020. doi:10.1016/j.neuron.2020.06.022.

[37] Johannes D Seelig and Vivek Jayaraman. Neural dynamics for landmark orientation and angular path integration. Nature, 521(7551):186–191, 2015.

[38] Floris van Breugel. A Nonlinear Observability Analysis of Ambient Wind Estimation with Uncalibrated Sensors, Inspired by Insect Neural Encoding. In 2021 60th IEEE Conference on Decision and Control (CDC), pages 1399–1406. IEEE, ec 2021. doi:10.1109/CDC45484.2021.9683219.

[39] D. C. O’Carroll, N. J. Bidweii, S. B. Laughlin, and E. J. Warrant. Insect motion detectors matched to visual ecology. Nature, 382(6586):63–66, July 1996. doi:10.1038/382063a0.

[40] Gaby Maimon, Andrew D Straw, and Michael H Dickinson. Active flight increases the gain of visual motion processing in drosophila. Nature Neuroscience, 13(3): 393–399, February 2010. doi:10.1038/nn.2492.

[41] Floris van Breugel, Renan Jewell, and Jaleesa Houle. Active anemosensing hypothesis: how flying insects could estimate ambient wind direction through sensory integration and active movement. Journal of The Royal Society Interface, 19(193):2022.03.31.486300, aug 2022. doi:10.1098/rsif.2022.0258.

[42] Floris van Breugel and Michael H. Dickinson. Plume-Tracking Behavior of Flying Drosophila Emerges from a Set of Distinct Sensory-Motor Reflexes. Current Biology, 24(3):274–286, feb 2014. doi:10.1016/j.cub.2013.12.023.

[43] Sawyer Buckminster Fuller, Andrew D. Straw, Martin Y. Peek, et al. Flying Drosophila stabilize their vision-based velocity controller by sensing wind with their antennae. Proceedings of the National Academy of Sciences, 111(13):E1182–E1191, apr 2014. doi:10.1073/pnas.1323529111.

[44] Debojyoti Biswas, Andrew Lamperski, Yu Yang, et al. Mode switching in organisms for solving explore-versus-exploit problems. Nature Machine Intelligence, 2023. doi:10.1038/s42256-023-00745-y.

[45] J. Humberto Ramos, Davis W. Adams, Kevin M. Brink, and Manoranjan Majji. Observability Informed Partial-Update Schmidt Kalman Filter. In 2021 IEEE 24th International Conference on Information Fusion (FUSION), pages 1–8. IEEE, nov 2021. doi:10.23919/FUSION49465.2021.9626946.

[46] Yvette E. Fisher, Michael Marquis, Isabel D’Alessandro, and Rachel I. Wilson. Dopamine promotes head direction plasticity during orienting movements. Nature, 612 (7939):316–322, dec 2022. doi:10.1038/s41586-022-05485-4.

[47] Roger A. Horn and Charles R. Johnson. Matrix Analysis. Cambridge University Press, oct 2012. doi:10.1017/CBO9781139020411.

[48] Y.H. Kim, F.L. Lewis, and C.T. Abdallah. Nonlinear observer design using dynamic recurrent neural networks. In Proceedings of 35th IEEE Conference on Decision and Control, volume 1, pages 949–954. IEEE, 1996. doi:10.1109/CDC.1996.574590.

[49] Young H. Kim, Frank L. Lewis, and Chaouki T. Abdallah. A dynamic recurrent neural-network-based adaptive observer for a class of nonlinear systems. Automatica, 33(8):1539–1543, aug 1997. doi:10.1016/S0005-1098(97)00065-4.

[50] S.N. Huang, K.K. Tan, and T.H. Lee. Further result on a dynamic recurrent neural-network-based adaptive observer for a class of nonlinear systems. Automatica, 41(12):2161–2162, ec 2005. doi:10.1016/j.automatica.2005.07.003.

[51] I. Salgado and I. Chairez. Discrete time recurrent neural network observer. In 2009 International Joint Conference on Neural Networks, pages 2764–2770. IEEE, jun 2009. doi:10.1109/IJCNN.2009.5178900.

[52] Benjamin Cellini and Jean-Michel Mongeau. Hybrid visual control in fly flight: insights into gaze shift via saccades. Current Opinion in Insect Science, 42:23–31, ec 2020. doi:10.1016/j.cois.2020.08.009.

[53] Bo. Cheng, S. N. Fry, Q. Huang, and X. Deng. Aerodynamic damping during rapid flight maneuvers in the fruit fly Drosophila. Journal of Experimental Biology, 213(4): 602–612, feb 2010. doi:10.1242/jeb.038778.

[54] F. van Breugel, Marie P. Suver, and Michael H. Dickinson. Octopaminergic modulation of the visual flight speed regulator of Drosophila. Journal of Experimental Biology, 217(10):1737–1744, may 2014. doi:10.1242/jeb.098665.

[55] Pauli Virtanen, Ralf Gommers, Travis E. Oliphant, et al. SciPy 1.0: fundamental algorithms for scientific computing in Python. Nature Methods, 17(3):261–272, mar 2020. doi:10.1038/s41592-019-0686-2.

[56] Felix Fiedler, Benjamin Karg, Lukas Lüken, et al. dompc: Towards FAIR nonlinear and robust model predictive control. Control Engineering Practice, 140:105676, nov 2023. doi:10.1016/j.conengprac.2023.105676.

[57] C Radhakrishna Rao, Sujit Kumar Mitra, et al. Generalized inverse of a matrix and its applications. In Proceedings of the sixth Berkeley symposium on mathematical statistics and probability, volume 1, pages 601–620. University of California Press Oakland, CA, USA, 1972.

[58] Martín Abadi, Paul Barham, Jianmin Chen, et al. TensorFlow: A system for large-scale machine learning. may 2016. doi:10.48550/arXiv.1605.08695.

[59] François and others Chollet. Keras, 2015.

